# A Lipid Prodrug Strategy Enhances Targeted Protein Degrader CNS Pharmacokinetics

**DOI:** 10.1101/2025.04.18.649560

**Authors:** Georges A. Leconte, Gillian E. Gadbois, Yadira Sepulveda, Ady Schwarz, Erin Chang, Raymond T. Suhandynata, Jeremiah D. Momper, Fleur M. Ferguson

## Abstract

Targeted protein degraders that recruit the von-hippel Lindau E3-ligase complex often have poor physicochemical properties, requiring extensive medicinal chemistry optimization prior to use in hypothesis-testing experiments *in vivo*, and with few blood-brain-barrier permeable examples disclosed. In this study, we systematically examine a panel of fatty acid promoieties as agents to enhance degrader pharmacokinetics and BBB-exposure. We characterize effects on cellular E3-ligase engagement, cellular BRD4 degradation kinetics, murine plasma stability, and murine blood plasma and brain pharmacokinetics and pharmacodynamics. We identify degrader prodrugs with significantly improved CNS exposure relative to the parent degrader. This led to successful BRD4. degradation in perfused brain samples, demonstrating that fatty acid promoieties can accelerate progress towards proof of principle *in vivo* experiments for CNS degrader projects.

## INTRODUCTION

Targeted protein degraders are powerful tools for facilitating chemical biology studies, target validation studies, and drug discovery efforts.^1^ However, PROTACs that recruit the von-hippel Lindau (VHL) E3-ligase complex tend to have less favourable physicochemical properties than those that recruit CRL4^CRBN^.^2^ This is due to the peptide-derived VHL-binding ligand, which is higher molecular weight, harbours a greater number of H-bond donors and acceptors, and has a larger topological polar surface area, compared to CRBN-recruiting gluteramide ligands.^3^ These effects can be exacerbated by the properties of polyethylene glycol (PEG) linkers often used in early-stage PROTACs to enhance flexibility and solubility.^4^ Consequently, early-stage VHL-recruiting degraders often require extensive medicinal chemistry optimization prior to use in hypothesis-testing experiments *in vivo*, and engineering brain exposure is a challenge. This resource-intensive step limits their accessibility as chemical biology tools, and delays critical target validation efforts.

Fatty acid and sterol functionalization approaches have been successfully used as a strategy to improve pharmacokinetic properties and brain exposure of drugs at the pre-clinical stage, such as the cholesterol functionalization of DNA/RNA heteroduplex antisense oligonucleotide therapies, which enabled CNS-exposure and target knockdown in murine brain.^5^ These promoieties are also found in FDA-approved drugs, such as Aripiprazole laurioxil, a longlasting injectable treatment for schitzophrenia, and Sobetirome-ethanolamine, where promoiety conjugation doubles the concentration of sobetirome achieved in patient brain (Fig. 1).^**6**, **7**^ Despite these successes, fatty acid promoiety conjugation has yet to be systematically applied to deliver heterobifunctional degraders across the BBB, which have distinct chemical properties from nucleotides and small molecule inhibitors.

**Figure 1.**
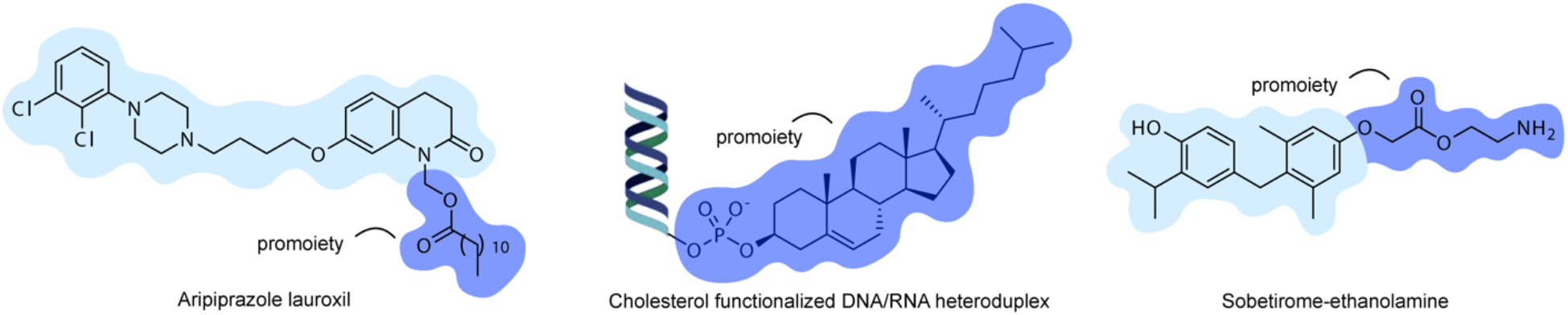
Examples of clinical and preclinical therapeutics where BBB permeability is enhanced by lipid and sterol promoieties. ^*5*, *8*, *9*^. Aripiprazole lauroxil is a long lasting injectable for treatment of schizophrenia in adults, while the parent drug is a daily oral treatment^10^. Cholesterol conjugation enabled RNA/DNA heteroduplexes to cross the BBB and knockdown genes in rodent CNS.^5^ Sobetirome-ethanolamine prodrug doubles sobetirome concentration in brain, and results in a more than 2-fold improved brain/serum ratio.^9^

In this work, we develop a library of prodrugs based on BRD4 degrader tools, and use them to explore the effects on the degraders cellular permeability, cellular promoiety release kinetics, cellular BRD4 degradation kinetics, and murine plasma stability. We perform snapshot murine blood plasma and brain pharmacokinetics and pharmacodynamics, and demonstrate enhanced brain exposure and degradation kinetics resulting from promoiety conjugation. We define key success parameters for enhancing BBB-permeability via lipidation, and a generalizable workflow and fatty acid library that can be applied to any VHL-recruiting degrader-of-interest.

## RESULTS

To design a promoiety library, we selected lipophilic biomolecules encompassing a range of fatty acyl lipid chain lengths from caproic acid (C6:0) to stearic acid (C18:0) (Fig. 2). We also selected tert-butyl esters for inclusion based on reports that tert-butyl esterification of CRBN-recruiting CDK2/4/6 PROTACs improved their oral bioavailability in mice.^11^ These promoieties were attached to the 4-*cis*-hydroxy-L-proline alcohol of VHL ligand 2^12^ via esterification, and incorporated into BRD4 degrader 5 (GAL-02-221, scheme 1).^13^ Functionalization at this position blocks ligand binding to VHL, until the promoiety is cleaved to release the free hydroxyl group. As lipidation can negatively impact the solubility of small molecules, all compounds were evaluated for solubility in DMSO, PBS, and cell culture media and found to have solubility sufficient to support testing at concentrations up to 100 µM.

**Figure 2.**
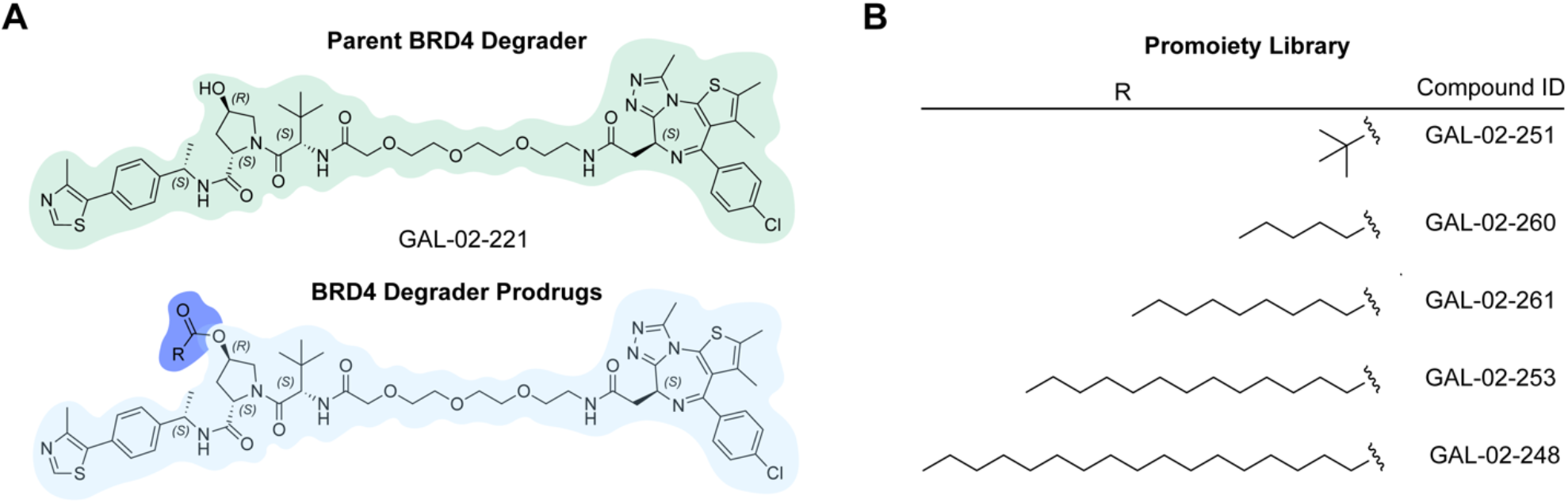
AproPROTAC library as a model for VHL-recruiting PROTACs. A. Structures of BRD4 degrader and prodrugs. B. Promoieties evaluated in this manuscript.

To quantify the proportion of active BRD4 degrader released into the cell, and evaluate the relative cellular promoiety release rates for each prodrug, we used a cellular VHL NanoBRET target engagement assay in HEK293 cells (Fig. 3A).^14^ In this assay, are the released VHL-recruiting degraders compete for binding to NanoLuc-tagged VHL with a BRET-acceptor-conjugated VHL ligand, reducing the BRET signal, while the promoiety conjugated degraders cannot bind to VHL and do not compete with the tracer. We selected 30 minute and 2 hour pre-incubation time points to evaluate the kinetics of intracellular prodrug release by esterases (Fig. 3C). Unmodified VHL ligand, and the parent BRD4 degrader GAL-02-221 were used as controls at each time point. As expected, these controls demonstrated negligible time-dependence in their IC_50_ values, indicating that they can rapidly permeate the cell to engage VHL (Fig. 3B-C).

**Figure 3.**
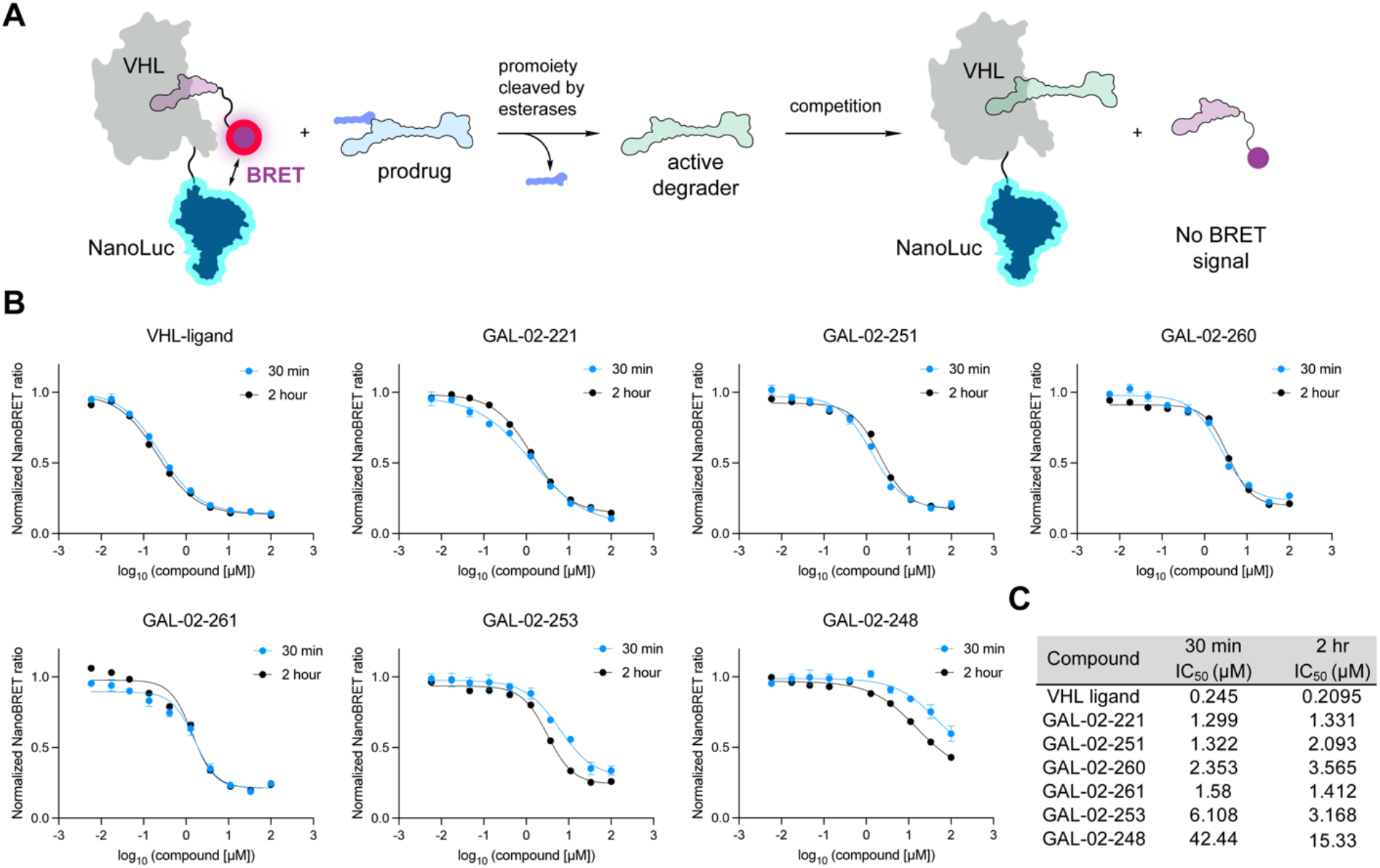
Time course VHL target engagement NanoBRET assay for measuring intracellular prodrug activation. A) A schematic representation of the assay wherein an active PROTAC (green), released from proPROTAC (blue), out competes a VHL ligand-BRET probe resulting in loss of BRET. B) Time-course IC_50_ data of parent VHL-binder, parent degrader and prodrugs in VHL-cellular target engagement assay. HEK293 cells transfected with VHL-NLuc were treated with the indicated concentration of pro**drug** and 1 µM tracer for the indicated time, followed by readout of BRET. Data presented as the average ± S.D. of *n = 3* replicates. C) Summary of IC_50_ data at 30 min and 2 hrs.

Parent degrader GAL-02-221 is approximately 5-fold less potent than VHL ligand in the VHL-target engagement assay, reflecting differences in their cell permeability (Fig. 3B-C). Tert-butyl ester GAL-02-251 and capric acid (C10:0) ester GAL-02-261 had comparable VHL engagement potency to unmodified GAL-02-221 at the 30 minute time point, indicating rapid promoiety cleavage and comparable cell permeability. Caproic acid (C6:0) conjugated prodrug GAL-02-260 had modestly reduced engagement of VHL that did not display time dependence, indicating reduced cell permeability relative to parent degrader GAL-02-221.

Prodrugs conjugated to the longer fatty acids myristic acid (C14:0, GAL-02-253) and stearic acid (C18:0, GAL-02-248) displayed lower VHL IC_50_s, and clear time-dependent improvements in target engagement, characteristic of a reduced intracellular cleavage rate of the promoiety by esterases. Together, this data demonstrated that all prodrugs were cell permeable, and that a range of intracellular promoiety release rates were represented in the library.

Next, we examined how the release kinetics of the prodrugs impacted their degradation kinetics, by measuring the intracellular activity of the prodrugs in a BRD4 HiBiT assay in Jurkat cells over a time course.^15^ In this assay, endogenous BRD4 is tagged at the C-terminus with a HiBiT peptide using CRISPR/cas9. Upon cell lysis and addition of LgBiT and a luciferin substrate, a functional NLuc is formed and luminescence is emitted proportional to BRD4-HiBiT abundance.^16^ Degradation was evaluated following 1 hr, 2 hr and 5 hr compound incubation times, which enabled us to evaluate changes in the degradation of BRD4 over time while to minimizing cell toxicity (Fig. 4).

**Figure 4.**
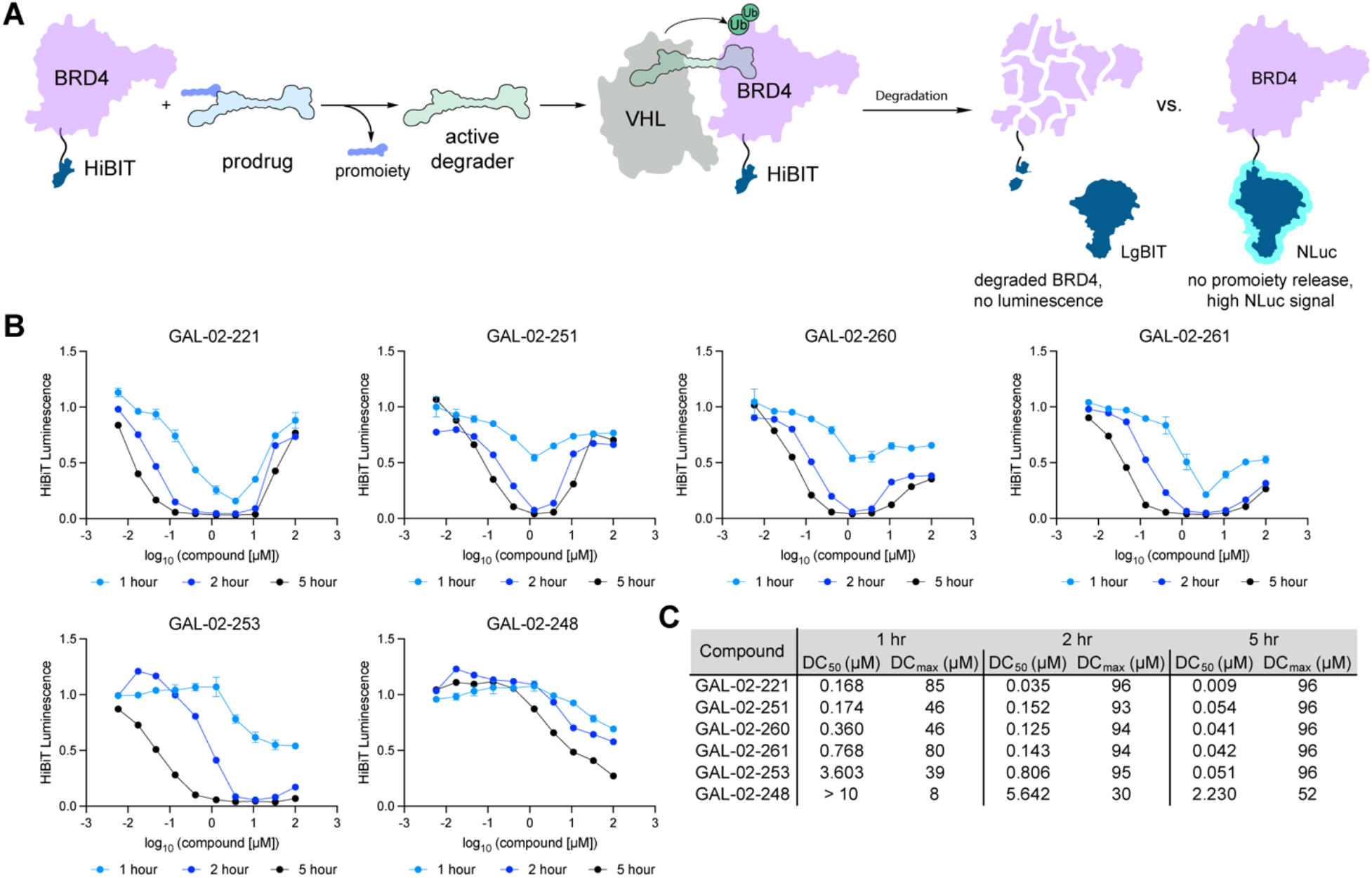
BRD4-HiBiT degradation assay. Time-course IC_50_ data of parent VHL-binder, parent degrader and prodrugs in VHL-cellular target engagement assay. A schematic representation of the BRD-HiBiT assay where upon release of prodrug (blue) to active degrader (green) in cells, BRD4-HiBiT is degraded. B). Time-course DC_50_ data of parent degrader and prodrugs. BRD4-HiBiT Jurkat cells were treated with the indicated concentration of proPROTAC for the indicated time. Data normalized to DMSO control and shown as mean +/-S.D. of *n=3* replicates. C) Summary of DC_50_ and D_max_ data depicted in B.

Following 1 hr incubation times, tert-butyl (GAL-02-251) and caproic acid (C6:0, GAL-02-260) modified analogues had significantly reduced D_max, 1hr_ compared to GAL-02-221, indicating that although complete prodrug release was observed within 30 minutes in the VHL-target engagement assay, the difference in concentrations of active GAL-02-221 during that initial time period has profound effects on cellular degradation kinetics. Capric acid prodrug (GAL-02-261, C10:0) showed comparable D_max, 1hr_ to GAL-02-221, but less potent DC_50,_ indicating promoiety release occurs faster than GAL-02-251 and GAL-01-260. Myristic acid (C14:0, GAL-02-253) and stearic acid (C18:0, GAL-02-248) conjugated analogues, whose promoieties were released more slowly, had reduced potency, and D_max, 1hr_ of 40% and 8 % respectively. By 2 hrs, GAL-02-251, GAL-02-260 and GAL-02-261 demonstrated comparable degradation of BRD4. Myristic acid (C14:0) analogue GAL-02-253 showed considerably increased degradation potency and depth by 2 hrs (D_max,2 hr_ 95%) and at 5 hrs, demonstrated comparable degradation potency as GAL-02-251, GAL-02-260 and GAL-02-261. The stearic acid prodrug (C18:0, GAL-02-248) remained weakly active at 2 hr and 5 hr time points consistent with reduced cellular promoiety release and reduced cell permeability.

As esterases are also present in blood plasma, we next sought to evaluate promoiety release kinetics in murine plasma (Fig. 5A-H). We performed plasma stability assays, treating murine plasma with 10 µM of each molecule and quantifying the levels of both prodrug and released parent degrader over a time course of 2 hrs by LC/MS/MS (Figure 5A-H, Table S1), with parent degrader GAL-02-221 included as a control. We observed robust plasma stability of parent compound GAL-02-221 over the assay, with 51% remaining at the assay end point (120 minutes, Fig. 5B). Prodrugs with short carbon chain promoieties (GAL-02-260, GAL-02-261) were rapidly cleaved in plasma, while tert-butyl and C14 analogues GAL-02-251 and GAL-02-253 showed moderate release kinetics. The longer C18 chain analogue GAL-02-248 demonstrated the slowest release kinetics, lowest signal, and incomplete recovery of GAL-02-221 at the 2 hrs time point. This indicates that GAL-02-248 has both a slower release rate and reduced availability compared to other prodrugs, consistent with the cellular assay data.

**Figure 5.**
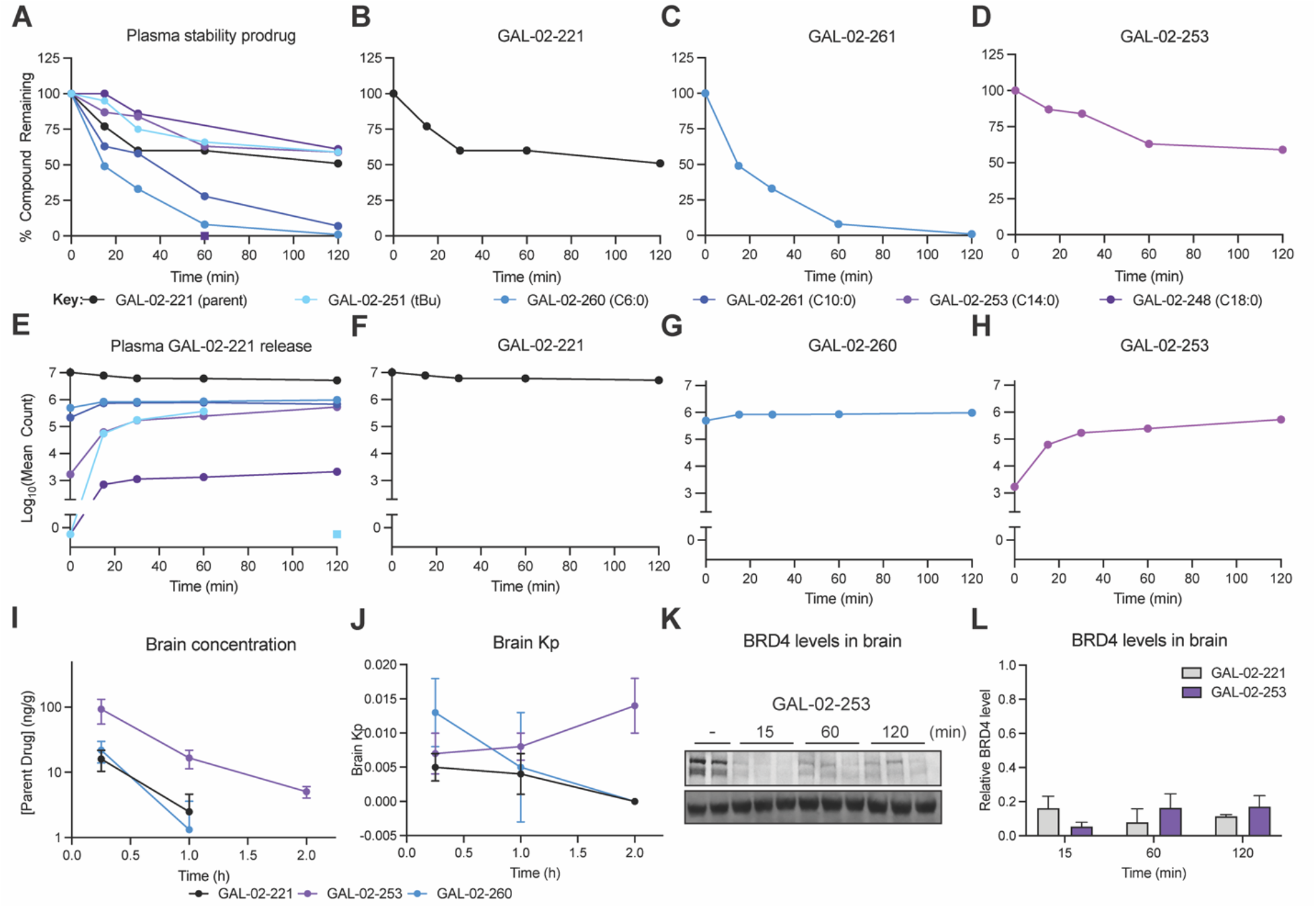
Prodrug stability and degradation kinetics in murine plasma and *in vivo*. A-D) *In vitro* murine plasma stability of prodrugs, concentration at time zero set to 100%, and E-H) Concentration of the released parent drug GAL-02-221 over 2 h. Values reported represent the mean of two independent experiments. F) *In vivo* brain concentration of the released parent drug GAL-02-221 over a 2 h period following a single IV injection (10 mpk) of the indicated compound. J) Brain Kp from the same experiment K) Western blot showing pharmacodynamic effect of the indicated compounds on BRD4 levels in perfused murine brain tissue. L) Quantification of K. Values reported represent the mean +/-S.D. of *n = 3* replicates.

Having characterized the cell and plasma pharmacology of the prodrugs, we next sought to evaluate the effect of lipidation on pharmacokinetic properties and CNS exposure. We selected GAL-02-253 and GAL-02-260 for snapshot pharmacokinetics, based on their cell permeability, activity, and differentiated promoiety release rates (fast, moderate). We performed intravenous dosing of C57BL/6 mice (*n =* 3 per compound) with 10 mg/kg (IV) of GAL-02-221 (reference degrader). These doses were adjusted to 12 mg/kg (IV) GAL-02-253 and 11 mg/kg (IV) GAL-02-260 to ensure an equivalent molarity of GAL-02-221 upon promoiety cleavage in all conditions. A relatively high dose was chosen to ensure accurate quantification by LC/MS/MS.

While the rapidly released GAL-02-260 prodrug had comparable pharmacokinetics to GAL-02-221, C14 conjugated degrader prodrug GAL-02-253 resulted in significantly increased GAL-02-221 exposure in plasma and brain (Fig. 5I-J, Fig. S3, Table S2). To assess if the improved exposure was sufficient to effect BRD4 degradation we performed pharmacodynamic studies on perfused brain and liver tissues following a single IV injection (Fig. 5K-L). Here, we observed potent degradation in liver by all compounds (Fig. S3). In brain, GAL-02-253 promoiety conjugation enabled superior rapid and complete BRD4 degradation at early time points, relative to GAL-02-221, though both were able to effect BRD4 degradation at these high doses (Fig. 5K-L, Fig. S3).

## DISCUSSION

Targeted protein degraders are infrequently applied to validate CNS targets due to the resource intensive medicinal chemistry optimization required to generate molecules that can enter and degrader proteins in the brain following systemic administration routes. In this work, we demonstrate that fatty acid promoieties can fast-track early stage degrader probes to *in vivo* experiments by enhancing their concentration in the CNS by up to 6-fold. While in our study we use high doses, to facilitate accurate quantification, leading to pharmacodynamic effects observed for both the free degrader and prodrug conjugated degrader in brain, these doses are unlikely to be tolerated in efficacy studies. In that context, a 6-fold improvement in brain exposure is highly significant as it may enabling longer-term experiments at tolerated doses. Targeted protein degraders are especially suited to these prodrug approaches, as their event-driven pharmacology leads to pharmacodynamic effects dependent on the peak concentration in brain C_max_, which we show is effectively modulated by the promoiety conjugation. This is in contrast to inhibitors, whose pharmacodynamic effects are dependent on the sustained exposure at efficacious concentrations in brain, which requires slower clearance (Cl_int_). More generally, we show that systematic evaluation of fatty acid promoieties can be applied to enable early-stage evaluation of VHL-recruiting degraders *in vivo*. We demonstrate that degrader palmitic acid prodrugs with which have moderate promoiety release kinetics (∼ 2hrs), and high cell permeability and activity at the 5 hrs time point are optimal for enhancing BBB-transport of degraders *in vivo*, providing a blueprint to guide broad application of this strategy across degrader programs.

Recent work has demonstrated that palmitic acid-derived promoieties can enhance cancer cell permeability of degraders via hijacking of the CD36 fatty acid transport receptor, *in vitro* and *in vivo*.^17^ Endogenously, CD36 is also expressed at the brain microvascular endothelia and functions to transport lipids across the blood brain barrier.^18^ In a disease context, loss of CD36 function enhances blood brain barrier function and is neuroprotective in Alzheimers Disease^18^ and Ischemic stroke^19^, suggesting CD36 may function as a permissive molecular transporter for facilitating BBB-passage beyond lipid scavenging. Here, we demonstrate that palimitic acid promoieties enhance BBB-permeability and brain exposure of degraders, *in vivo*, addressing a large unmet need in preclinical degrader development for neuroscience applications. We hypothesize that both CD36 mediated active transport and caveolae-mediated transcytosis are possible mediators of the enhanced BB-transport. Future work is needed to interrogate the precise mechanism of BBB-transport for the degraders and promoiety conjugates, in the context of intact and disrupted BBB.

## Supporting information

Biology methods

Chemistry methods

## AUTHOR CONTRIBUTIONS

GAL synthesized the molecules with assistance from AS and EC. GEG performed cellular VHL target engagement and BRD4 degradation assays. YS performed plasma stability assays. JM and RS performed study design of plasma stability experiments. FMF and GS performed data interpretation of PK experiments. FMF conceived the study, supervised the work and wrote the manuscript, with edits from all authors.

## ACKNOWLEDGEMENTS

This work was funded by a PhRMA Foundation Start-up Grant (FMF and JM, 30149761), NIH DP2NS132610 (FMF), an ACS BRIDGE Fellowship (GAL), NIH Molecular Biophysics Training Grant T32GM139795 (GEG), UCSD Undergraduate Summer Research Fellowship (AS, EC). We would like to thank J.J. LaClair for assistance with preparation of the TOC figure. The BRD4 HiBiT Jurkat cell line was a gift from N.S. Gray and I. You. We thank JJ LaClair for assistance with illustrations.

## TOC IMAGE

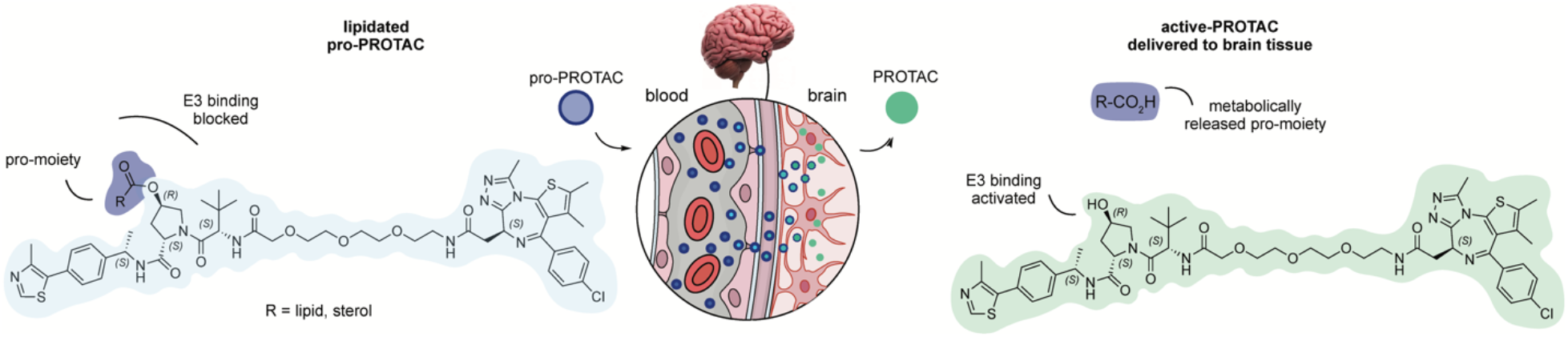

## Supporting Figures

**Figure S1.**
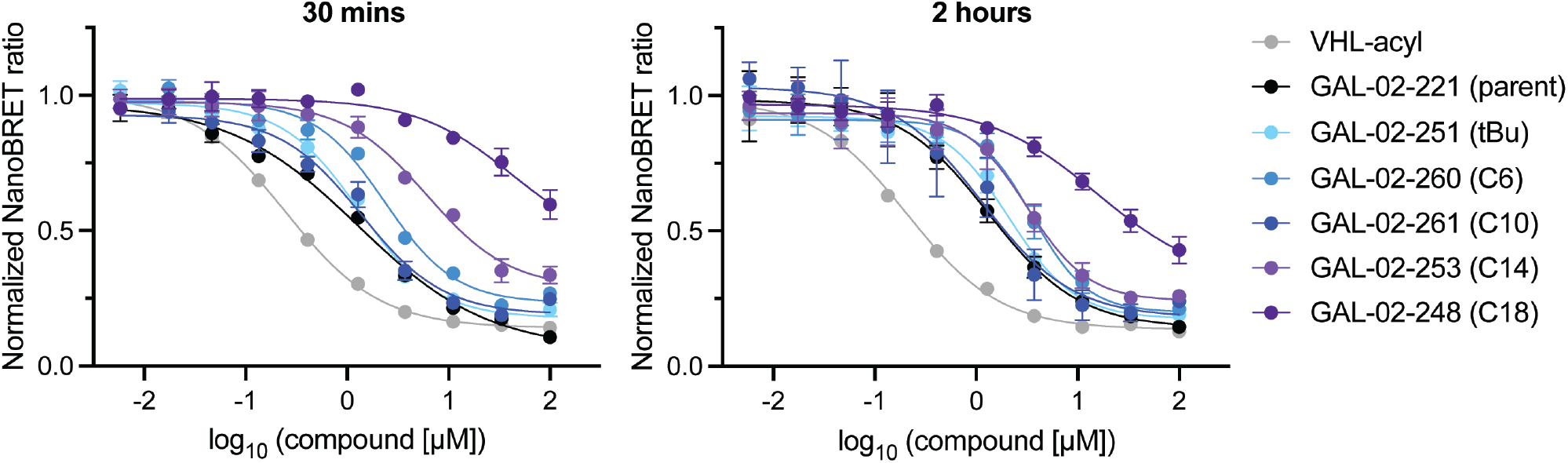
Time dependent VHL target engagement by proPROTACs. HEK293 cells expressing VHL-NLuc were treated with 1µM tracer and indicated concentration of proPROTAC for 30 minutes or 2 hours. BRET donor and emission signals were measured following the addition of the 3X Complete Substrate Plus Inhibitor Solution. The data was background corrected by subtracting BRET signal in the absence of tracer and then normalized to DMSO BRET signal. Data shown as the average of n = 3 replicates +/-SD.

**Figure S2.**
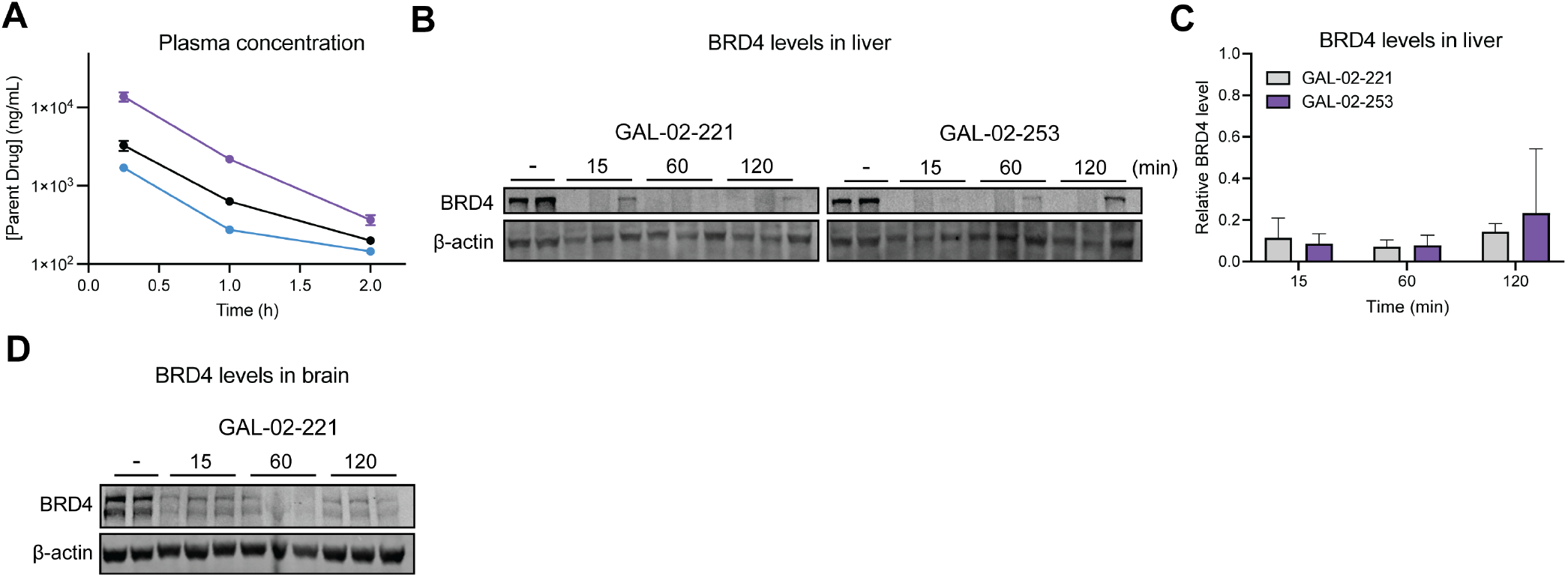
Pharmacodynamic effect in liver of degrader prodrugs. A) Snapshot pharmacokinetics of proPROTACs, measuring exposure of GAL-02-221 in plasma over 2 hour time course. B) Western blot of BRD4 in murine liver tissue following treatment with either GAL-02-221 or GAL-02-253 for 15, 60 or 120 minutes. C) Quantification of B. Values represent the mean +/-S.D. of *n = 3* replicates. D) Western blot of BRD4 in murine brain tissue following treatment with GAL-02-221 for 15, 60 or 120 minutes.

**Figure S3.**
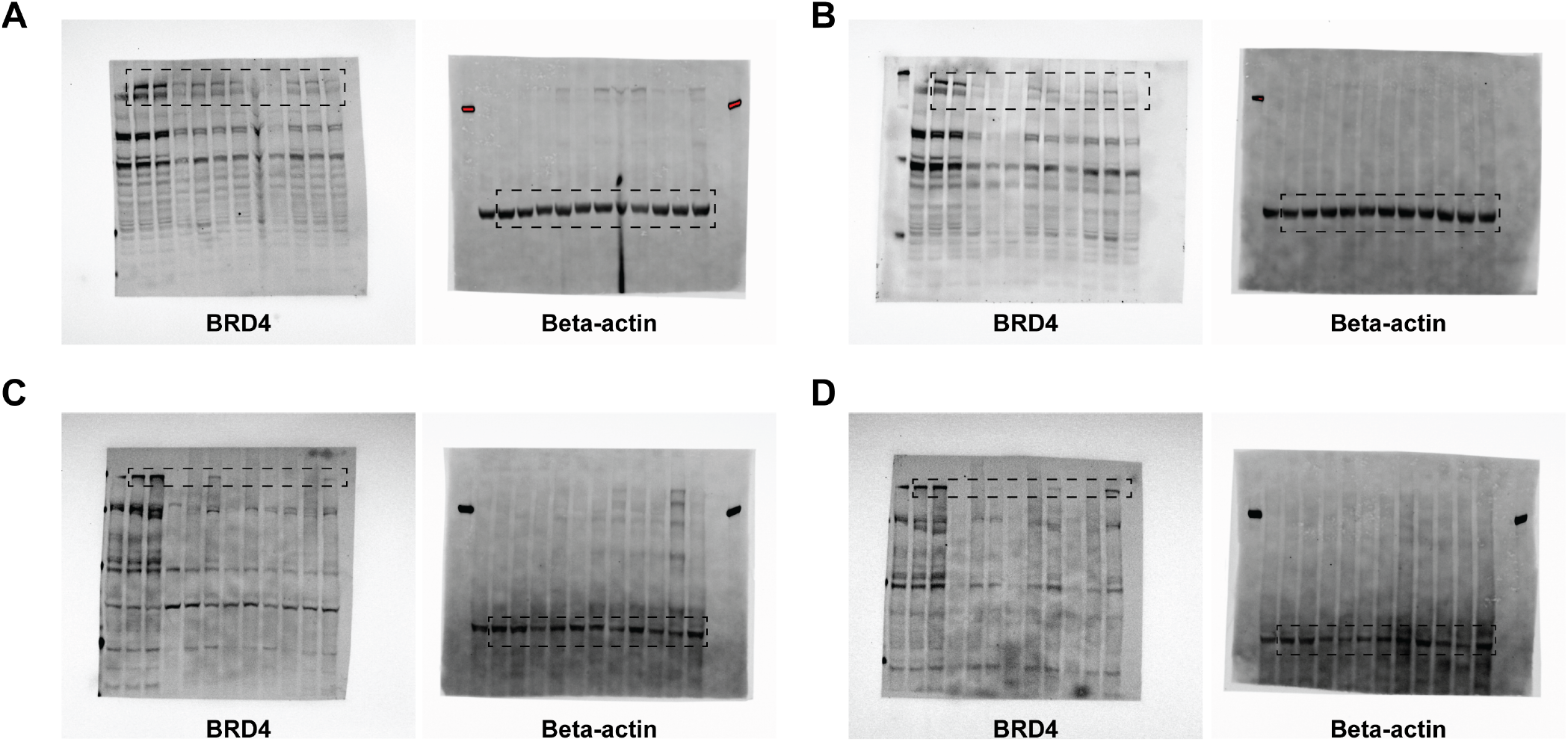
Uncropped western blots. A) Uncropped western blot of GAL-02-221 treated murine brain tissue. B) Uncropped western blot of GAL-02-253 treated murine brain tissue. C) Uncropped western blot of GAL-02-221 treated murine liver tissue. D) Uncropped western blot of GAL-02-253 treated murine liver tissue.

