## Supplementary material for "A Lipid Prodrug Strategy Enhances Targeted Protein Degrader CNS Pharmacokinetics": Biology methods

**Cell culture**
HEK293 (ATCC, CRL-1573) were cultured in high-glucose DMEM (Gibco, 11965092) with 10% FBS (Gibco, 10437028) and 1% Penicillin–Streptomycin (Gibco, 15140122). Jurkat BRD4-HiBiT cells were cultured in RPMI-1640 media (Gibco, 11875093) with 10% FBS (Gibco, 10437028) and 1% Penicillin–Streptomycin (Gibco, 15140122). All cells were maintained in 37 °C and 5% CO2 incubators and routinely tested negative for mycoplasma contamination using the MycoAlert Kit (Lonza, LT07318).

**VHL Target engagement assay**
The Promega VHL NanoBRET target engagement assay (Promega, N2931) was used to quantify VHL target engagement of the compounds. HEK293 cells were counted and adjusted to 200,000 cells/mL in Opti-MEM (Gibco, 11058021) with 1% FBS and transfected with VHL-NanoLuc® construct. The transfection mixture was prepared by mixing 9.0µg/mL Transfection Carrier DNA, 1.0µg/mL of VHL-NanoLuc® fusion vector DNA and TransIT-2020 (Mirus, 5400) in Opti-MEM and incubating mixture for 20 minutes. Cells and transfection mixture were combined at a 20:1 cell to transfection complex ratio. The cells were then transferred to a T75 flask and incubated for 24 hr at 37°C, 5% CO2. The following day, 34µL of cells at 200,000 cells/mL in Opti-MEM were plated into 384-well white, nonbinding surface microplates (Corning, 3574) and co-treated in triplicate with 1µM tracer and a 10-point half-log titration of compound at 100µM in DMSO using a Multidrop Pico 8 Digital Dispenser (Thermo). The plates were then incubated at 37°C, 5% CO2 for desired time point. Plates were allowed to equilibrate to room temperature for 15 min prior to characterization. Then 17µL of 3X Complete Substrate Plus Inhibitor Solution was added to each well of the 384-well microplate. Plates were incubated for 2 min at room temperature and then donor emission wavelength (450nm) and acceptor emission wavelength (610nm) were measured using a ClarioSTAR Plus microplate reader (BMG Labtech). To calculate BRET ratios the acceptor signal/emission signal for the no-tracer background control was subtracted from the acceptor signal/emission signal for the treatment condition and then that value was multiplied by 1000. These values were then normalized to the DMSO BRET ratios. Data was plotted in GraphPad Prism and EC50 calculated using a log(inhibitor) vs response – variable slope.

**BRD4-HiBiT degradation assay**
Jurkat BRD4-HiBiT cells were counted and adjusted to 200,000 cells/mL in complete RMPI media. 50µL of cell suspension was plated into each well of a white 384-well microplate (Corning, 3570). Multidrop Pico 8 digital dispenser (Thermo) was used to dose the cells with a 10-point half-log titration of compound starting at 100µM in DMSO in triplicate The plates were then incubated at 37°C, 5% CO2 for desired time point. Promega HiBiT lytic detection kit (Promega, N3030) was used to determine the HiBiT signal. 25µL of the Nano-Glo HiBiT lytic reagent was added to each well and the plate was shaken on an orbital shaker for 10 minutes at 600rpm. Luminescence was measured using ClarioSTAR Plus microplate reader (BMG Labtech). The raw data was DMSO normalized, log-transformed and plotted in GraphPad Prism.

**Plasma Stability**
Plasma stability of compounds were evaluated by the addition of the study compounds to preheated (37°C) mouse plasma to yield a final concentration of 10µM of study compound. Plasma stability assays were performed in a circular rotating incubator at 37°C in duplicate. 10µl aliquots were taken at 0 (immediately after addition of study compounds), 15, 30, 60, and 120 min and added to 30µl of cold (4°C) protein crash solution (acetonitrile, 0.1% formic acid) to deproteinize the plasma. The samples were mixed by vortexing for 1 min and then centrifugation at 4°C for 15 min at 10,000 rpm. The clear supernatants were analyzed by LC-MS/MS on an API

4000 triple quadrupole mass spectrometer (AB SCIEX). Values reported represent the mean of two independent experiments.

**Pharmacokinetic studies**

*Animals:* C57BL/6 mice (male, 8-12 weeks, 20-24g) were housed in standard laboratory conditions, with food and water provided ad libitum. Animals were dosed intravenously with the indicated compound (200µL).

*Compound Formulation:* 5% N-methylpyrrolidone, 5 % Solutol HS-15 and 90% Normal Saline.

*Sample collection and analysis.* Approximately 250 µL blood was collected under light isoflurane anesthesia (Surgivet®) through retro-orbital sinus from a set of three mice at 0.25, 1 and 2 hr post-dose followed by collection of CSF. Immediately after blood collection samples were treated with 1 mM PMSF, plasma was harvested by centrifugation at 10000 rpm, 10 min at 4ºC and samples were stored at -70±10ºC until bioanalysis. Following whole blood and CSF collection, animals were anesthetized and the whole body was perfused using 10 mL of sterile PBS pH 7.4. Brain, liver and spleen samples were collected from a set of three mice at 0.25, 1 and 2 h. After isolation, brain samples were rinsed three times in ice cold PBS (for 5-10 seconds/rinsed using ~5-10 mL PBS in a disposable petri dish for each rinse) dried on blotting paper and weighed. Each brain sample was divided into two equal parts and half of the brain was submitted for bioanalysis and the other half, along with spleen and liver, were flash frozen in LN2 and stored for future PD study. All the samples were stored below -70±10 ºC until bioanalysis. Note: Brain samples were diluted (1-part tissue: 2-parts of PBS) and homogenized. The homogenate was submitted for bioanalysis and the concentrations (ng/mL) received were corrected for dilution factor (3-fold) and the final reported concentrations were represented in ng/g. Samples were analyzed by LC-MS/MS on an API 4000 triple quadruple mass spectrometer.

*Data analysis.* Pharmacokinetic parameters for each test compound were calculated using standard non-compartmental approaches (Phoenix WinNonlin software Version 8.3).

**Western blot**

Mice tissue were lysed with ice cold RIPA buffer containing 1X HALT inhibitor cocktail (Thermo, 78442) and 125 U/mLbenzonase (MilliporeSigma, 706643). The samples were sonicated for 2 minutes with 10 sec on 3 sec off pulses at 70% power (Qsonica, Q125). Lysates were clarified by centrifugation at 21,000 x *g* for 15 minutes at 4 °C. BCA assay (Thermo, 23225) was performed according to manufacturer instructions to determine total protein concentration. SDS-PAGE samples were run on Bolt 4-12% Bis-Tris Gels (Invitrogen, NW04125BOX) in MES run buffer for 1 hour at 180 V. Gels were transferred to nitrocellulose membranes (Cytiva, 10600011) in Bolt transfer buffer (Invitrogen, BT00061) for 90 min at 45 V. Membranes were blocked with 5% non-fat dry milk (Kroger) in TBST (Thermo, 28360) for 1 hour at room temperature. Membranes were incubated with 1:10,000 BRD4 (Bethyl, A301-985A50) and 1:1,000 beta-actin (Cell Signaling Technology, 3700S) primary antibody diluted in TBS blocking buffer (LI-COR, 92760001) overnight at 4 °C. Membranes were washed three times with TBST then incubated with DyLight 680 anti-mouse IgG (Cell Signaling Technology, 5470S) and DyLight 800 antirabbit IgG (Cell Signaling Technology, 5151S) diluted 1:10,000 in TBS blocking buffer for 1 hour at room temperature. Following incubation with secondary antibodies, membranes were washed with TBST three times then imaged with a ChemiDoc Imaging System (Bio-Rad) and quantified in Image Lab 6.1.0 (Bio-Rad).
