## Supplementary material for "A Lipid Prodrug Strategy Enhances Targeted Protein Degrader CNS Pharmacokinetics": Chemistry methods

### Synthetic Methods

#### General Procedures

Unless otherwise noted, reagents and solvents were obtained from commercial suppliers and were used without further purification. $^1$H NMR spectra were recorded on 500 MHz Bruker Avance III spectrometer or 500 MHz JEOL ECA 500 spectrometer and chemical shifts are reported in parts per million (ppm, δ) downfield from tetramethylsilane (TMS). Coupling constants (J) are reported in Hz. Spin multiplicities are described as s (singlet), br (broad singlet), d (doublet), t (triplet), q (quartet), and m (multiplet). Mass spectra were obtained on a Waters Acquity UPLC/MS. Preparative HPLC was performed on a Waters Sunfire C18 column (19 mm × 50 mm, 5 μM) using a gradient of 15−95% methanol in water containing 0.05% trifluoroacetic acid (TFA) over 22 min (28 min run time) at a flow rate of 20 mL/min. Assayed compounds were isolated and tested as TFA salts. Purities of assayed compounds were in all cases greater than 95%, as determined by reverse-phase HPLC analysis.

#### Procedure A for the Deprotection of Tert-butoxyl carbamates

**tert-butyl ((S)-1-((2S,4R)-4-hydroxy-2-(((S)-1-(4-(4-methylthiazol-5-yl)phenyl)ethyl)carbamoyl)pyrrolidin-1-yl)-3,3-dimethyl-1-oxobutan-2-yl)carbamate** (1.0eq) was dissolved in 4N HCl in dioxane (5.0eq) and stirred at room temperature until completion was observed by LC-MS. At completion, the reaction was concentrated under reduced pressure and then dried on vacuum at 40°C for 1.5h to yield **(2S,4R)-1-((S)-2-amino-3,3-dimethylbutanoyl)-4-hydroxy-N-((S)-1-(4-(4-methylthiazol-5-yl)phenyl)ethyl)pyrrolidine-2-carboxamide hydrochloride** as a pale solid used without further purification.

#### Procedure B for the synthesis of VHL-linker analogues

**(2S,4R)-1-((S)-2-amino-3,3-dimethylbutanoyl)-4-hydroxy-N-((S)-1-(4-(4-methylthiazol-5-yl)phenyl)ethyl)pyrrolidine-2-carboxamide hydrochloride** (1.0eq), 2,2-dimethyl-4-oxo-3,8,11,14-tetraoxa-5-azahexadecan-16-oic acid (1.2eq), HATU (2.0eq), were dissolved in with DMF (5mL) in a flask charged with a stir bar. DIPEA (5.0eq) was added dropwise and the reaction was allowed to stir at room temperature for 18hr. At completion, the reaction was diluted with ethyl acetate and transferred to a separatory funnel where it was extracted with saturated sodium bicarbonate (2 x 10mL), 5% citric acid (2 x 10mL), and brine (1x 15mL). The resultant organic layer was dried over $MgSO4_4$ and purified via flash chromatography. Relevant fractions were identified, combined, and concentrated under reduced pressure to yield **tert-butyl ((S)-13-((2S,4R)-4-hydroxy-2-(((S)-1-(4-(4-methylthiazol-5-yl)phenyl)ethyl)carbamoyl)pyrrolidine-1-carbonyl)-14,14-dimethyl-11-oxo-3,6,9-trioxa-12-azapentadecyl)carbamate** as a pale yellow solid.

#### Procedure C for the synthesis of GAL-02-221 Analogues

**(2S,4R)-1-((S)-14-amino-2-(tert-butyl)-4-oxo-6,9,12-trioxa-3-azatetradecanoyl)-4-hydroxy-N-((S)-1-(4-(4-methylthiazol-5-yl)phenyl)ethyl)pyrrolidine-2-carboxamide hydrochloride** (1.0eq), (S)-2-(4-(4-chlorophenyl)-2,3,9-trimethyl-6H-thieno[3,2-f][1,2,4]triazolo[4,3-a][1,4]diazepin-6-yl)acetic acid (1.2eq) and HATU (2.0eq) were dissolved in DMF (5mL) in a flask charged with a stir bar. DIPEA (5.0eq) was added dropwise and the reaction was allowed to stir at room temperature for 18hr. At completion, the reaction was diluted with ethyl acetate and transferred to a separatory funnel where it was extracted with saturated sodium bicarbonate (2 x

10mL), 5% citric acid (2 x 10mL), and brine (1x 15mL). The resultant organic layer was dried over $MgSO4_4$ and purified via preparatory HPLC. Relevant fractions were identified, combined, and lyophilized to yield **(2S,4R)-1-((S)-2-(tert-butyl)-17-((S)-4-(4-chlorophenyl)-2,3,9-trimethyl-6H-thieno[3,2-f][1,2,4]triazolo[4,3-a][1,4]diazepin-6-yl)-4,16-dioxo-6,9,12-trioxa-3,15-diazaheptadecanoyl)-4-hydroxy-N-((S)-1-(4-(4-methylthiazol-5-yl)phenyl)ethyl)pyrrolidine-2-carboxamide** as a pale lyophilized solid.

#### Procedure D for the Synthesis of VHL Promoieties

**tert-butyl ((S)-1-((2S,4R)-4-hydroxy-2-(((S)-1-(4-(4-methylthiazol-5-yl)phenyl)ethyl)carbamoyl)pyrrolidin-1-yl)-3,3-dimethyl-1-oxobutan-2-yl)carbamate** (1.0eq) was added to a flask charged with a stir bar, dissolved in dichloromethane, cooled to 0 ° C on ice, and stirred vigorously until dissolved. To the stirring solution, triethylamine (4.0eq) was added and allowed to incubate at 0 °C for 5 minutes. Following, the pivaloyl chloride (1.5eq) dissolved in dichloromethane was added to the vigorously stirring reaction dropwise, after which the reaction was allowed to warm slowly to room temperature and stir for two hours. At completion, the reaction mixture was diluted with more dichloromethane and transferred to a separatory funnel and partitioned with saturated sodium bicarbonate where it was extracted twice. The resultant organic layer was dried over $MgSO_4$ and purified via flash chromatography. Relevant fractions were identified, combined, and concentrated under reduced pressure to yield **(3R,5S)-1-((S)-2-((tert-butoxycarbonyl)amino)-3,3-dimethylbutanoyl)-5-(((S)-1-(4-(4-methylthiazol-5-yl)phenyl)ethyl)carbamoyl)pyrrolidin-3-yl pivalate as a pale solid**.

#### Synthesis of GAL-02-221

**(2S,4R)-1-((S)-2-amino-3,3-dimethylbutanoyl)-4-hydroxy-N-((S)-1-(4-(4-methylthiazol-5-yl)phenyl)ethyl)pyrrolidine-2-carboxamide hydrochloride (1)**

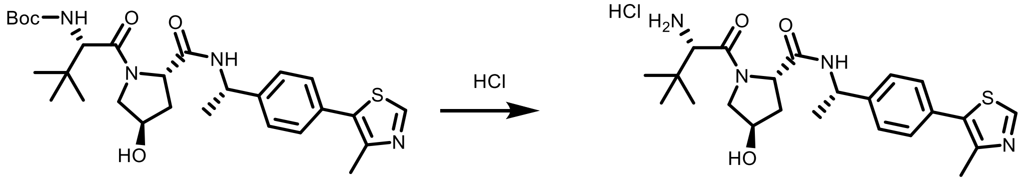


(2S,4R)-1-((S)-2-amino-3,3-dimethylbutanoyl)-4-hydroxy-N-((S)-1-(4-(4-methylthiazol-5-yl)phenyl)ethyl)pyrrolidine-2-carboxamide hydrochloride (quantitative) was synthesized according to general procedure A and used without further purification. MS (ESI) m/z 444 $[M+H]^+$

**tert-butyl ((S)-13-((2S,4R)-4-hydroxy-2-(((S)-1-(4-(4-methylthiazol-5-yl)phenyl)ethyl)carbamoyl)pyrrolidine-1-carbonyl)-14,14-dimethyl-11-oxo-3,6,9-trioxa-12-azapentadecyl)carbamate**

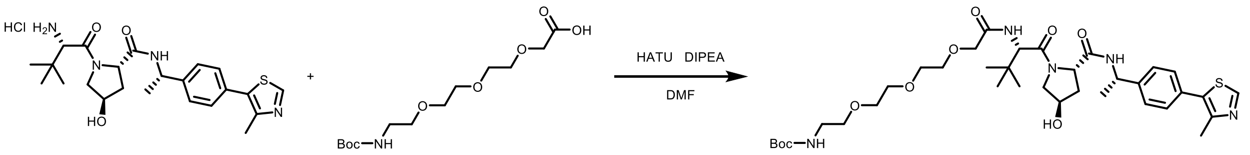


tert-butyl ((S)-13-((2S,4R)-4-hydroxy-2-(((S)-1-(4-(4-methylthiazol-5-yl)phenyl)ethyl)carbamoyl)pyrrolidine-1-carbonyl)-14,14-dimethyl-11-oxo-3,6,9-trioxa-12-

azapentadecyl)carbamate (571mg, 100% yield) was synthesized according to general procedure B. MS (ESI) m/z 634 $[M+H-100]^+$

**(2S,4R)-1-((S)-14-amino-2-(tert-butyl)-4-oxo-6,9,12-trioxa-3-azatetradecanoyl)-4-hydroxy-N-((S)-1-(4-(4-methylthiazol-5-yl)phenyl)ethyl)pyrrolidine-2-carboxamide hydrochloride**

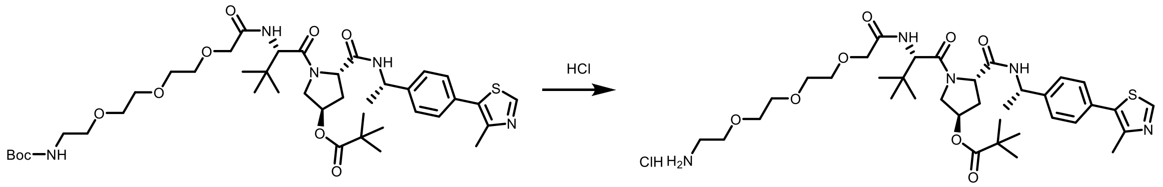


(2S,4R)-1-((S)-14-amino-2-(tert-butyl)-4-oxo-6,9,12-trioxa-3-azatetradecanoyl)-4-hydroxy-N-((S)-1-(4-(4-methylthiazol-5-yl)phenyl)ethyl)pyrrolidine-2-carboxamide hydrochloride (quantitative) was synthesized according to general procedure A. MS (ESI) m/z 634 $[M+H]^+$

**(2S,4R)-1-((S)-2-(tert-butyl)-17-((S)-4-(4-chlorophenyl)-2,3,9-trimethyl-6H-thieno[3,2-f][1,2,4]triazolo[4,3-a][1,4]diazepin-6-yl)-4,16-dioxo-6,9,12-trioxa-3,15-diazaheptadecanoyl)-4-hydroxy-N-((S)-1-(4-(4-methylthiazol-5-yl)phenyl)ethyl)pyrrolidine-2-carboxamide**

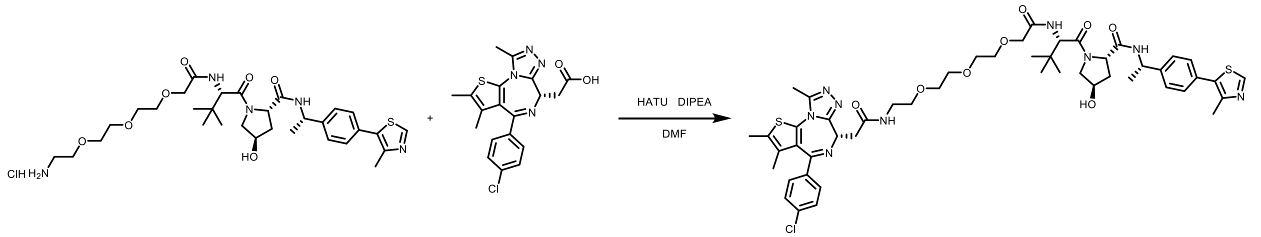


(2S,4R)-1-((S)-2-(tert-butyl)-17-((S)-4-(4-chlorophenyl)-2,3,9-trimethyl-6H-thieno[3,2-f][1,2,4]triazolo[4,3-a][1,4]diazepin-6-yl)-4,16-dioxo-6,9,12-trioxa-3,15-diazaheptadecanoyl)-4-hydroxy-N-((S)-1-(4-(4-methylthiazol-5-yl)phenyl)ethyl)pyrrolidine-2-carboxamide (17mg, 10% yield) was synthesized according to general procedure C. MS (ESI) m/z 1015. $^1$H NMR (500 MHz, DMSO-D6) δ 8.99 (s, 1H), 8.46 (d, J = 7.7 Hz, 1H), 8.32 (t, J = 5.7 Hz, 1H), 7.48 (d, J = 8.3 Hz, 2H), 7.44 – 7.41 (m, 4H), 7.36 (d, J = 8.2 Hz, 2H), 4.90 (t, J = 7.2 Hz, 1H), 4.56 – 4.49 (m, 2H), 4.44 (t, J = 8.1 Hz, 1H), 4.27 (s, 1H), 3.97 (s, 2H), 3.61 (dd, J = 4.8, 2.5 Hz, 2H), 3.57 (td, J = 8.6, 4.1 Hz, 8H), 3.46 (t, J = 5.9 Hz, 2H), 3.31 – 3.25 (m, 2H), 2.60 (s, 3H), 2.45 (s, 3H), 2.40 (s, 3H), 2.07 – 2.00 (m, 1H), 1.76 (ddd, J = 13.1, 8.9, 4.5 Hz, 1H), 1.61 (s, 3H), 1.36 (d, J = 7.1 Hz, 3H), 0.93 (s, 9H).

### Synthesis of ProPROTACs

**(3R,5S)-1-((S)-2-((tert-butoxycarbonyl)amino)-3,3-dimethylbutanoyl)-5-(((S)-1-(4-(4-methylthiazol-5-yl)phenyl)ethyl)carbamoyl)pyrrolidin-3-yl pivalate**

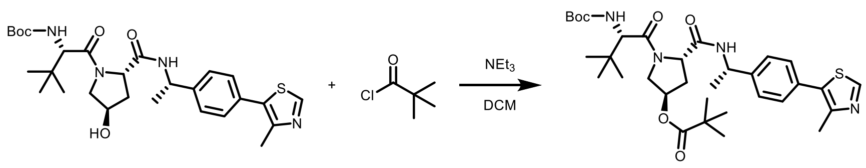

x

(3R,5S)-1-((S)-2-((tert-butoxycarbonyl)amino)-3,3-dimethylbutanoyl)-5-(((S)-1-(4-(4-methylthiazol-5-yl)phenyl)ethyl)carbamoyl)pyrrolidin-3-yl pivalate (231mg, 61% yield) was synthesized according to general procedure D. MS (ESI) m/z 529 [M+H-100]$^+$

**(3R,5S)-1-((S)-2-((tert-butoxycarbonyl)amino)-3,3-dimethylbutanoyl)-5-(((S)-1-(4-(4-methylthiazol-5-yl)phenyl)ethyl)carbamoyl)pyrrolidin-3-yl hexanoate**

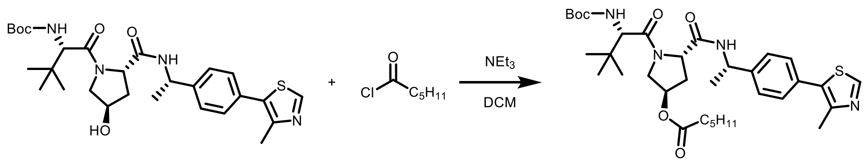

x

(3R,5S)-1-((S)-2-((tert-butoxycarbonyl)amino)-3,3-dimethylbutanoyl)-5-(((S)-1-(4-(4-methylthiazol-5-yl)phenyl)ethyl)carbamoyl)pyrrolidin-3-yl hexanoate (243mg, 98% yield) was synthesized according to general procedure D. MS (ESI) m/z 543 [M+H-100]$^+$

**(3R,5S)-1-((S)-2-((tert-butoxycarbonyl)amino)-3,3-dimethylbutanoyl)-5-(((S)-1-(4-(4-methylthiazol-5-yl)phenyl)ethyl)carbamoyl)pyrrolidin-3-yl decanoate**

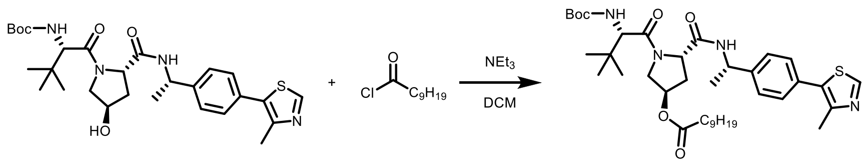

x

(3R,5S)-1-((S)-2-((tert-butoxycarbonyl)amino)-3,3-dimethylbutanoyl)-5-(((S)-1-(4-(4-methylthiazol-5-yl)phenyl)ethyl)carbamoyl)pyrrolidin-3-yl decanoate (222mg, 82% yield) was synthesized according to general procedure D. MS (ESI) m/z 599 [M+H-100]$^+$

**(3R,5S)-1-((S)-2-((tert-butoxycarbonyl)amino)-3,3-dimethylbutanoyl)-5-(((S)-1-(4-(4-methylthiazol-5-yl)phenyl)ethyl)carbamoyl)pyrrolidin-3-yl tetradecanoate**

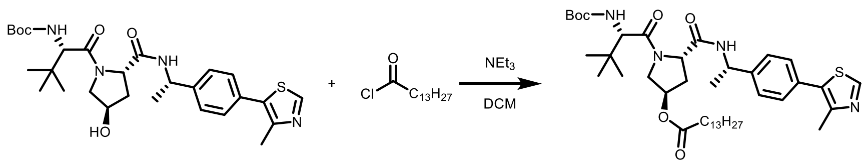

x

(3R,5S)-1-((S)-2-((tert-butoxycarbonyl)amino)-3,3-dimethylbutanoyl)-5-(((S)-1-(4-(4-methylthiazol-5-yl)phenyl)ethyl)carbamoyl)pyrrolidin-3-yl tetradecanoate (516mg, 93% yield) was synthesized according to general procedure D. MS (ESI) m/z 655 [M+H-100]$^+$

**(3R,5S)-1-((S)-2-((tert-butoxycarbonyl)amino)-3,3-dimethylbutanoyl)-5-(((S)-1-(4-(4-methylthiazol-5-yl)phenyl)ethyl)carbamoyl)pyrrolidin-3-yl stearate**

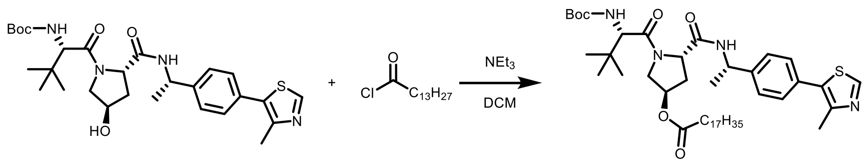


x

(3R,5S)-1-((S)-2-((tert-butoxycarbonyl)amino)-3,3-dimethylbutanoyl)-5-(((S)-1-(4-(4-methylthiazol-5-yl)phenyl)ethyl)carbamoyl)pyrrolidin-3-yl stearate (202mg, 90% yield) was synthesized according to general procedure D. MS (ESI) m/z 711 [M+H-100]

**(3R,5S)-1-((S)-2-amino-3,3-dimethylbutanoyl)-5-(((S)-1-(4-(4-methylthiazol-5-yl)phenyl)ethyl)carbamoyl)pyrrolidin-3-yl pivalate hydrochloride**

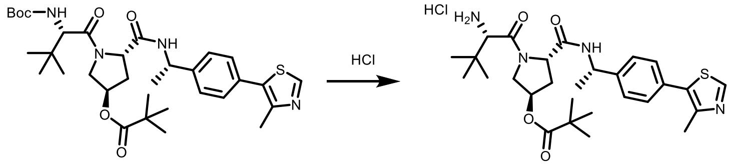


(3R,5S)-1-((S)-2-amino-3,3-dimethylbutanoyl)-5-(((S)-1-(4-(4-methylthiazol-5-yl)phenyl)ethyl)carbamoyl)pyrrolidin-3-yl pivalate hydrochloride (quantitative) was synthesized according to general procedure A. MS (ESI) m/z 529 $[M+H]^+$

**(3R,5S)-1-((S)-2-amino-3,3-dimethylbutanoyl)-5-(((S)-1-(4-(4-methylthiazol-5-yl)phenyl)ethyl)carbamoyl)pyrrolidin-3-yl hexanoate hydrochloride**

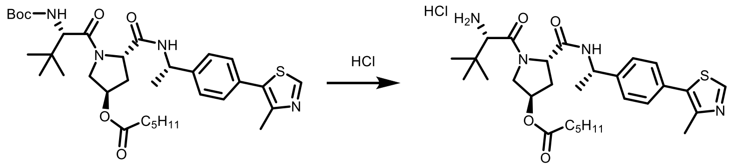


(3R,5S)-1-((S)-2-amino-3,3-dimethylbutanoyl)-5-(((S)-1-(4-(4-methylthiazol-5-yl)phenyl)ethyl)carbamoyl)pyrrolidin-3-yl hexanoate hydrochloride (quantitative) was synthesized according to general procedure A. MS (ESI) m/z 543 $[M+H]^+$

**(3R,5S)-1-((S)-2-amino-3,3-dimethylbutanoyl)-5-(((S)-1-(4-(4-methylthiazol-5-yl)phenyl)ethyl)carbamoyl)pyrrolidin-3-yl decanoate hydrochloride**

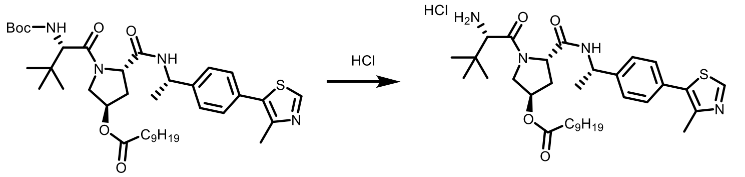


(3R,5S)-1-((S)-2-amino-3,3-dimethylbutanoyl)-5-(((S)-1-(4-(4-methylthiazol-5-yl)phenyl)ethyl)carbamoyl)pyrrolidin-3-yl decanoate hydrochloride (quantitative) was synthesized according to general procedure A. MS (ESI) m/z 599 $[M+H]^+$

**(3R,5S)-1-((S)-2-amino-3,3-dimethylbutanoyl)-5-(((S)-1-(4-(4-methylthiazol-5-yl)phenyl)ethyl)carbamoyl)pyrrolidin-3-yl tetradecanoate hydrochloride**

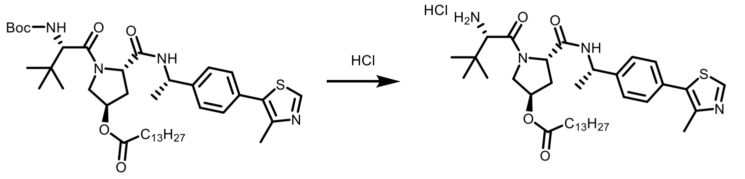


(3R,5S)-1-((S)-2-amino-3,3-dimethylbutanoyl)-5-(((S)-1-(4-(4-methylthiazol-5-yl)phenyl)ethyl)carbamoyl)pyrrolidin-3-yl tetradecanoate hydrochloride (quantitative) was synthesized according to general procedure A. MS (ESI) m/z 655 $[M+H]^+$

**(3R,5S)-1-((S)-2-amino-3,3-dimethylbutanoyl)-5-(((S)-1-(4-(4-methylthiazol-5-yl)phenyl)ethyl)carbamoyl)pyrrolidin-3-yl stearate hydrochloride**

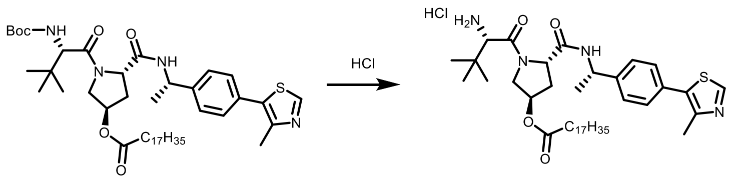


(3R,5S)-1-((S)-2-amino-3,3-dimethylbutanoyl)-5-(((S)-1-(4-(4-methylthiazol-5-yl)phenyl)ethyl)carbamoyl)pyrrolidin-3-yl stearate hydrochloride (quantitative) was synthesized according to general procedure A. m/z 711 $[M+H]^+$

**(3R,5S)-1-((S)-18-(tert-butyl)-2,2-dimethyl-4,16-dioxo-3,8,11,14-tetraoxa-5,17-diazanonadecan-19-oyl)-5-(((S)-1-(4-(4-methylthiazol-5-yl)phenyl)ethyl)carbamoyl)pyrrolidin-3-yl pivalate (ETC-01-013)**

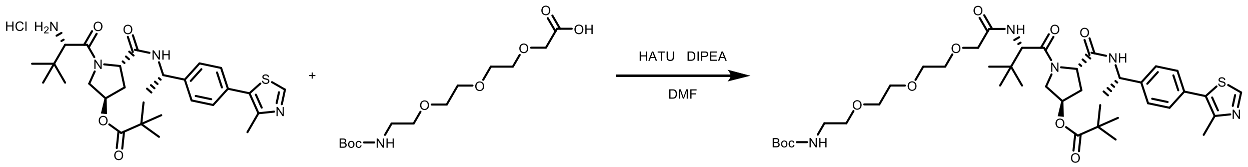


(3R,5S)-1-((S)-18-(tert-butyl)-2,2-dimethyl-4,16-dioxo-3,8,11,14-tetraoxa-5,17-diazanonadecan-19-oyl)-5-(((S)-1-(4-(4-methylthiazol-5-yl)phenyl)ethyl)carbamoyl)pyrrolidin-3-yl pivalate (94mg, 83% yield) was synthesized according to general procedure B. MS (ESI) m/z 718 $[M+H-100]^+$

**(3R,5S)-1-((S)-18-(tert-butyl)-2,2-dimethyl-4,16-dioxo-3,8,11,14-tetraoxa-5,17-diazanonadecan-19-oyl)-5-(((S)-1-(4-(4-methylthiazol-5-yl)phenyl)ethyl)carbamoyl)pyrrolidin-3-yl hexanoate**

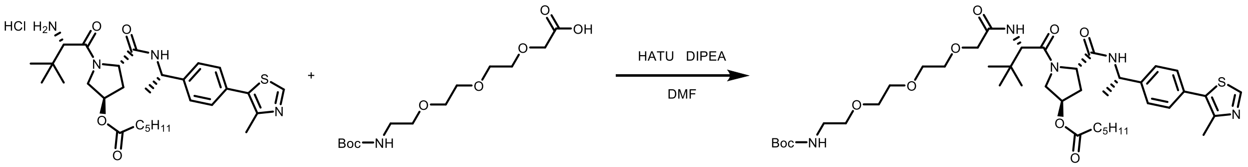


(3R,5S)-1-((S)-18-(tert-butyl)-2,2-dimethyl-4,16-dioxo-3,8,11,14-tetraoxa-5,17-diazanonadecan-19-oyl)-5-(((S)-1-(4-(4-methylthiazol-5-yl)phenyl)ethyl)carbamoyl)pyrrolidin-3-yl hexanoate

(104mg, 44% yield) was synthesized according to general procedure B. MS (ESI) m/z 732 [M+H-100]$^+$

**(3R,5S)-1-((S)-18-(tert-butyl)-2,2-dimethyl-4,16-dioxo-3,8,11,14-tetraoxa-5,17-diazanonadecan-19-oyl)-5-(((S)-1-(4-(4-methylthiazol-5-yl)phenyl)ethyl)carbamoyl)pyrrolidin-3-yl decanoate**

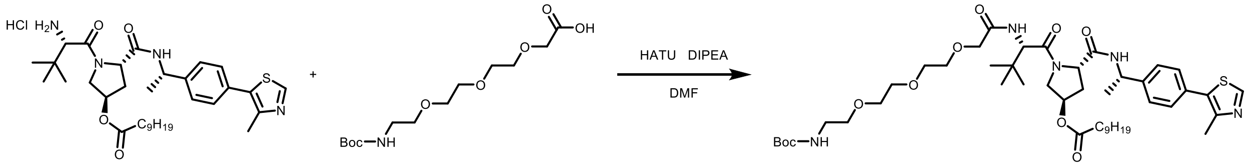


(3R,5S)-1-((S)-18-(tert-butyl)-2,2-dimethyl-4,16-dioxo-3,8,11,14-tetraoxa-5,17-diazanonadecan-19-oyl)-5-(((S)-1-(4-(4-methylthiazol-5-yl)phenyl)ethyl)carbamoyl)pyrrolidin-3-yl decanoate (126mg, 49% yield) was synthesized according to general procedure B. MS (ESI) m/z 788 [M+H-100]$^+$

**(3R,5S)-1-((S)-18-(tert-butyl)-2,2-dimethyl-4,16-dioxo-3,8,11,14-tetraoxa-5,17-diazanonadecan-19-oyl)-5-(((S)-1-(4-(4-methylthiazol-5-yl)phenyl)ethyl)carbamoyl)pyrrolidin-3-yl tetradecanoate**

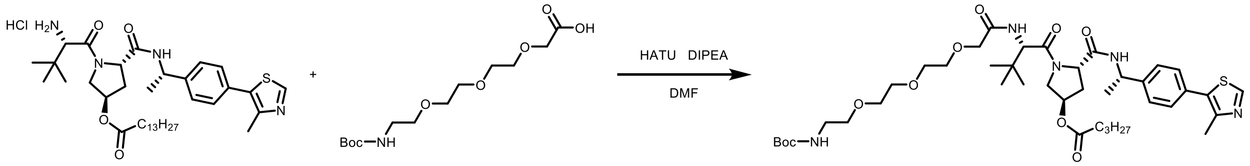


(3R,5S)-1-((S)-18-(tert-butyl)-2,2-dimethyl-4,16-dioxo-3,8,11,14-tetraoxa-5,17-diazanonadecan-19-oyl)-5-(((S)-1-(4-(4-methylthiazol-5-yl)phenyl)ethyl)carbamoyl)pyrrolidin-3-yl tetradecanoate (196mg, 32% yield) was synthesized according to general procedure B. MS (ESI) m/z 844 [M+H-100]$^+$

**(3R,5S)-1-((S)-18-(tert-butyl)-2,2-dimethyl-4,16-dioxo-3,8,11,14-tetraoxa-5,17-diazanonadecan-19-oyl)-5-(((S)-1-(4-(4-methylthiazol-5-yl)phenyl)ethyl)carbamoyl)pyrrolidin-3-yl stearate**

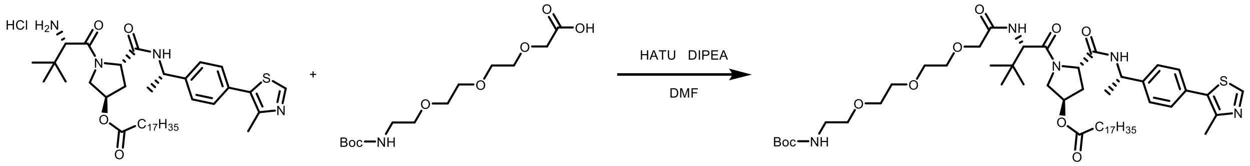


(3R,5S)-1-((S)-18-(tert-butyl)-2,2-dimethyl-4,16-dioxo-3,8,11,14-tetraoxa-5,17-diazanonadecan-19-oyl)-5-(((S)-1-(4-(4-methylthiazol-5-yl)phenyl)ethyl)carbamoyl)pyrrolidin-3-yl stearate (171mg, 85% yield) was synthesized according to general procedure B. MS (ESI) m/z 900 [M+H-100]$^+$

**(3R,5S)-1-((S)-14-amino-2-(tert-butyl)-4-oxo-6,9,12-trioxa-3-azatetradecanoyl)-5-(((S)-1-(4-(4-methylthiazol-5-yl)phenyl)ethyl)carbamoyl)pyrrolidin-3-yl pivalate hydrochloride**

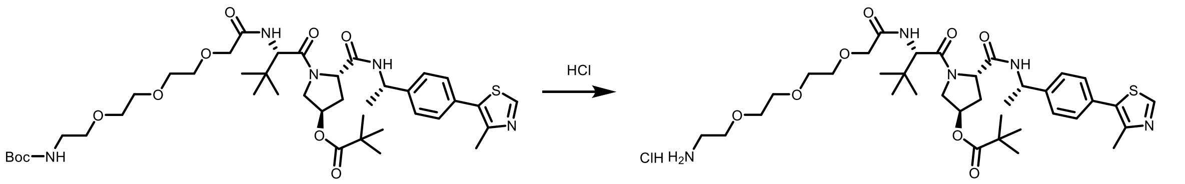


(3R,5S)-1-((S)-14-amino-2-(tert-butyl)-4-oxo-6,9,12-trioxa-3-azatetradecanoyl)-5-(((S)-1-(4-(4-methylthiazol-5-yl)phenyl)ethyl)carbamoyl)pyrrolidin-3-yl pivalate hydrochloride (quantitative) was synthesized according to general procedure A. MS (ESI) m/z 718 $[M+H]^+$

**(3R,5S)-1-((S)-14-amino-2-(tert-butyl)-4-oxo-6,9,12-trioxa-3-azatetradecanoyl)-5-(((S)-1-(4-(4-methylthiazol-5-yl)phenyl)ethyl)carbamoyl)pyrrolidin-3-yl hexanoate hydrochloride**

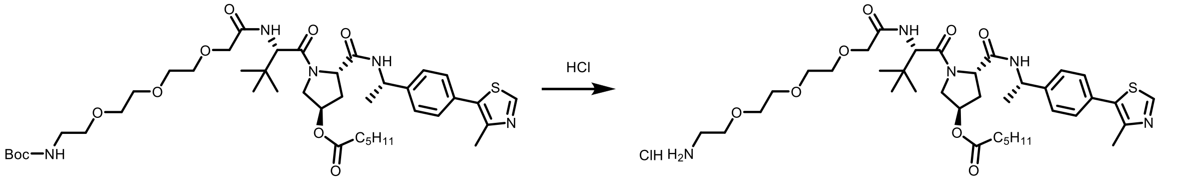


(3R,5S)-1-((S)-14-amino-2-(tert-butyl)-4-oxo-6,9,12-trioxa-3-azatetradecanoyl)-5-(((S)-1-(4-(4-methylthiazol-5-yl)phenyl)ethyl)carbamoyl)pyrrolidin-3-yl hexanoate hydrochloride (quantitative) was synthesized according to general procedure A. MS (ESI) m/z 732 $[M+H]^+$

**(3R,5S)-1-((S)-14-amino-2-(tert-butyl)-4-oxo-6,9,12-trioxa-3-azatetradecanoyl)-5-(((S)-1-(4-(4-methylthiazol-5-yl)phenyl)ethyl)carbamoyl)pyrrolidin-3-yl decanoate hydrochloride**

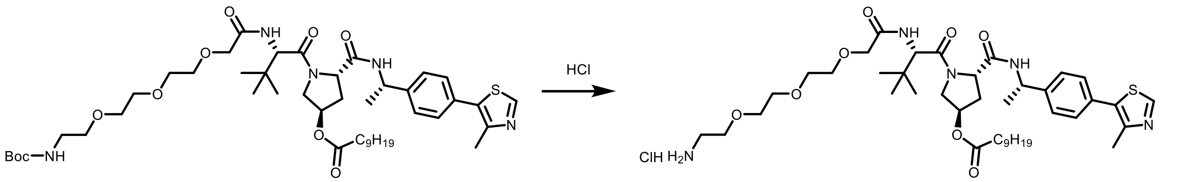


(3R,5S)-1-((S)-14-amino-2-(tert-butyl)-4-oxo-6,9,12-trioxa-3-azatetradecanoyl)-5-(((S)-1-(4-(4-methylthiazol-5-yl)phenyl)ethyl)carbamoyl)pyrrolidin-3-yl decanoate hydrochloride (quantitative) was synthesized according to general procedure A. MS (ESI) m/z 788 $[M+H]^+$

**(3R,5S)-1-((S)-14-amino-2-(tert-butyl)-4-oxo-6,9,12-trioxa-3-azatetradecanoyl)-5-(((S)-1-(4-(4-methylthiazol-5-yl)phenyl)ethyl)carbamoyl)pyrrolidin-3-yl tetradecanoate hydrochloride**

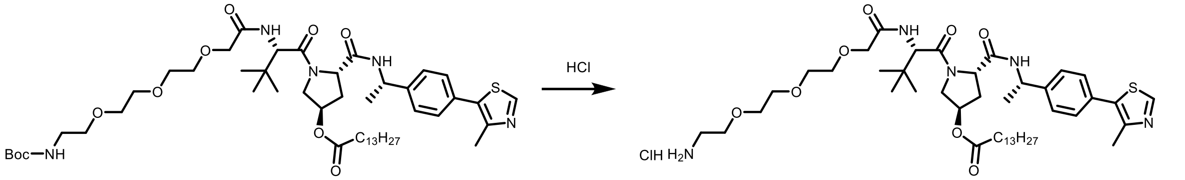


(3R,5S)-1-((S)-14-amino-2-(tert-butyl)-4-oxo-6,9,12-trioxa-3-azatetradecanoyl)-5-(((S)-1-(4-(4-methylthiazol-5-yl)phenyl)ethyl)carbamoyl)pyrrolidin-3-yl tetradecanoate hydrochloride (quantitative) was synthesized according to general procedure A. MS (ESI) m/z 844 $[M+H]^+$

**(3R,5S)-1-((S)-14-amino-2-(tert-butyl)-4-oxo-6,9,12-trioxa-3-azatetradecanoyl)-5-(((S)-1-(4-(4-methylthiazol-5-yl)phenyl)ethyl)carbamoyl)pyrrolidin-3-yl stearate hydrochloride**

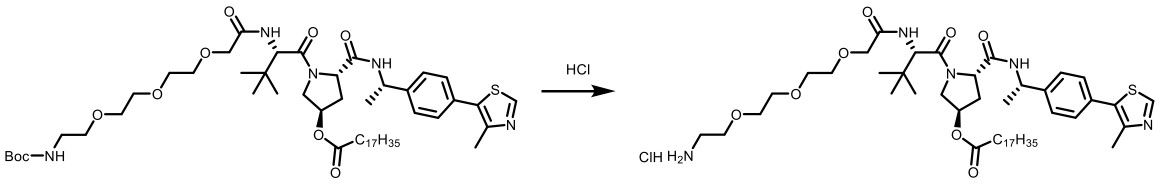


(3R,5S)-1-((S)-14-amino-2-(tert-butyl)-4-oxo-6,9,12-trioxa-3-azatetradecanoyl)-5-(((S)-1-(4-(4-methylthiazol-5-yl)phenyl)ethyl)carbamoyl)pyrrolidin-3-yl tetradecanoate hydrochloride (quantitative) was synthesized according to general procedure A. MS (ESI) m/z 900 $[M+H]^+$

**(3R,5S)-1-((S)-2-(tert-butyl)-17-((S)-4-(4-chlorophenyl)-2,3,9-trimethyl-6H-thieno[3,2-f][1,2,4]triazolo[4,3-a][1,4]diazepin-6-yl)-4,16-dioxo-6,9,12-trioxa-3,15-diazaheptadecanoyl)-5-(((S)-1-(4-(4-methylthiazol-5-yl)phenyl)ethyl)carbamoyl)pyrrolidin-3-yl pivalate (GAL-02-251)**

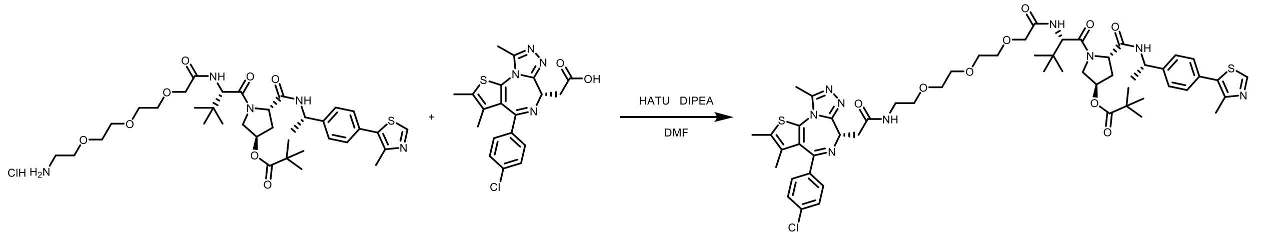


(3R,5S)-1-((S)-2-(tert-butyl)-17-((S)-4-(4-chlorophenyl)-2,3,9-trimethyl-6H-thieno[3,2-f][1,2,4]triazolo[4,3-a][1,4]diazepin-6-yl)-4,16-dioxo-6,9,12-trioxa-3,15-diazaheptadecanoyl)-5-(((S)-1-(4-(4-methylthiazol-5-yl)phenyl)ethyl)carbamoyl)pyrrolidin-3-yl pivalate (43mg, 31% yield) was synthesized according to general procedure C. MS (ESI) m/z 1099. $^1$H NMR (400 MHz, DMSO-D6) δ 8.90 (s, 1H), 8.42 (d, J = 7.7 Hz, 1H), 8.22 (t, J = 5.6 Hz, 1H), 7.41 – 7.37 (m, 2H), 7.35 – 7.31 (m, 4H), 7.30 – 7.25 (m, 3H), 5.07 (s, 1H), 4.81 (t, J = 7.2 Hz, 1H), 4.42 (dd, J = 7.8, 5.9 Hz, 1H), 4.39 – 4.34 (m, 2H), 3.84 (s, 2H), 3.81 – 3.80 (m, 2H), 3.25 – 3.08 (m, 4H), 2.50 (s, 3H), 2.41 (p, J = 1.8 Hz, 7H), 2.35 (s, 3H), 2.31 (s, 3H), 1.52 (s, 3H), 1.27 (d, J = 7.0 Hz, 2H), 0.99 (s, 8H), 0.85 (d, J = 3.6 Hz, 9H).

**(3R,5S)-1-((S)-2-(tert-butyl)-17-((S)-4-(4-chlorophenyl)-2,3,9-trimethyl-6H-thieno[3,2-f][1,2,4]triazolo[4,3-a][1,4]diazepin-6-yl)-4,16-dioxo-6,9,12-trioxa-3,15-diazaheptadecanoyl)-5-(((S)-1-(4-(4-methylthiazol-5-yl)phenyl)ethyl)carbamoyl)pyrrolidin-3-yl hexanoate (GAL-02-260)**

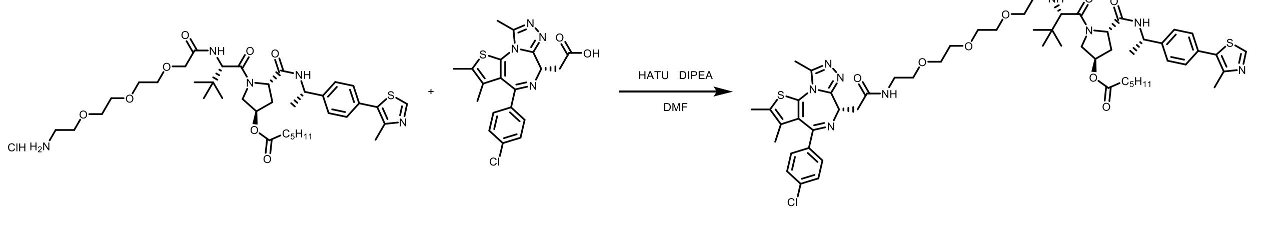


(3R,5S)-1-((S)-2-(tert-butyl)-17-((S)-4-(4-chlorophenyl)-2,3,9-trimethyl-6H-thieno[3,2-f][1,2,4]triazolo[4,3-a][1,4]diazepin-6-yl)-4,16-dioxo-6,9,12-trioxa-3,15-diazaheptadecanoyl)-5-(((S)-1-(4-(4-methylthiazol-5-yl)phenyl)ethyl)carbamoyl)pyrrolidin-3-yl hexanoate (4mg, 2% yield) was synthesized according to general procedure C. MS (ESI) m/z 1113. $^1$H NMR (500 MHz, DMSO-D6) δ 9.00 (s, 1H), 8.49 (d, J = 7.7 Hz, 1H), 8.32 (t, J = 5.7 Hz, 1H), 7.48 (d, J = 8.7 Hz,

2H), 7.43 (dd, J = 4.2, 1.9 Hz, 2H), 7.43 – 7.40 (m, 2H), 7.37 (t, J = 6.4 Hz, 3H), 5.20 (s, 1H), 4.94 – 4.85 (m, 1H), 4.51 (dd, J = 8.1, 6.1 Hz, 1H), 4.48 – 4.41 (m, 2H), 3.99 – 3.89 (m, 2H), 3.87 (d, J = 11.9 Hz, 1H), 3.63 – 3.54 (m, 8H), 3.46 (t, J = 5.9 Hz, 2H), 3.34 – 3.26 (m, 2H), 3.22 (td, J = 14.2, 5.3 Hz, 3H), 2.60 (s, 3H), 2.45 (s, 3H), 2.40 (s, 3H), 2.24 (dh, J = 14.5, 4.0 Hz, 3H), 1.97 (td, J = 9.0, 4.6 Hz, 1H), 1.61 (s, 3H), 1.48 (h, J = 6.7 Hz, 3H), 1.36 (d, J = 6.9 Hz, 2H), 1.22 (td, J = 10.8, 5.6 Hz, 4H), 0.94 (d, J = 7.5 Hz, 9H), 0.83 (t, J = 7.0 Hz, 3H).

**(3R,5S)-1-((S)-2-(tert-butyl)-17-((S)-4-(4-chlorophenyl)-2,3,9-trimethyl-6H-thieno[3,2-f][1,2,4]triazolo[4,3-a][1,4]diazepin-6-yl)-4,16-dioxo-6,9,12-trioxa-3,15-diazaheptadecanoyl)-5-(((S)-1-(4-(4-methylthiazol-5-yl)phenyl)ethyl)carbamoyl)pyrrolidin-3-yl decanoate (GAL-03-261)**

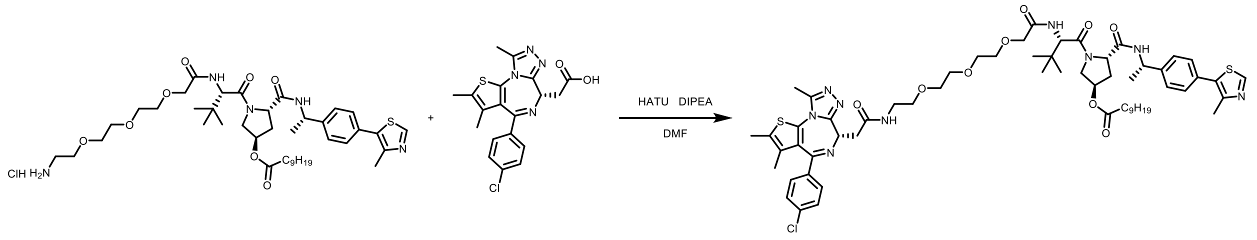


(3R,5S)-1-((S)-2-(tert-butyl)-17-((S)-4-(4-chlorophenyl)-2,3,9-trimethyl-6H-thieno[3,2-f][1,2,4]triazolo[4,3-a][1,4]diazepin-6-yl)-4,16-dioxo-6,9,12-trioxa-3,15-diazaheptadecanoyl)-5-(((S)-1-(4-(4-methylthiazol-5-yl)phenyl)ethyl)carbamoyl)pyrrolidin-3-yl decanoate (16mg, 8% yield) was synthesized according to general procedure C. MS (ESI) m/z 1169. $^1$H NMR (500 MHz, DMSO-D6) δ 8.96 (s, 1H), 8.45 (d, J = 7.7 Hz, 1H), 8.28 (t, J = 5.7 Hz, 1H), 7.45 (d, J = 8.7 Hz, 2H), 7.40 (d, J = 3.1 Hz, 2H), 7.39 – 7.36 (m, 2H), 7.34 (t, J = 6.4 Hz, 2H), 5.15 (d, J = 21.2 Hz, 1H), 4.87 (p, J = 7.1 Hz, 1H), 4.48 (dd, J = 8.1, 6.0 Hz, 1H), 4.45 – 4.37 (m, 2H), 3.95 – 3.87 (m, 2H), 3.84 (d, J = 11.8 Hz, 1H), 3.62 – 3.48 (m, 8H), 3.42 (t, J = 5.9 Hz, 2H), 3.32 – 3.13 (m, 3H), 2.56 (s, 3H), 2.42 (s, 3H), 2.37 (s, 3H), 2.21 (dh, J = 12.2, 4.0 Hz, 3H), 1.94 (ddd, J = 13.7, 9.2, 4.8 Hz, 1H), 1.58 (s, 3H), 1.49 – 1.38 (m, 2H), 1.33 (d, J = 7.0 Hz, 2H), 1.19 (s, 12H), 0.91 (d, J = 8.3 Hz, 9H), 0.80 (t, J = 6.9 Hz, 3H).

**(3R,5S)-1-((S)-2-(tert-butyl)-17-((S)-4-(4-chlorophenyl)-2,3,9-trimethyl-6H-thieno[3,2-f][1,2,4]triazolo[4,3-a][1,4]diazepin-6-yl)-4,16-dioxo-6,9,12-trioxa-3,15-diazaheptadecanoyl)-5-(((S)-1-(4-(4-methylthiazol-5-yl)phenyl)ethyl)carbamoyl)pyrrolidin-3-yl tetradecanoate (GAL-02-253)**

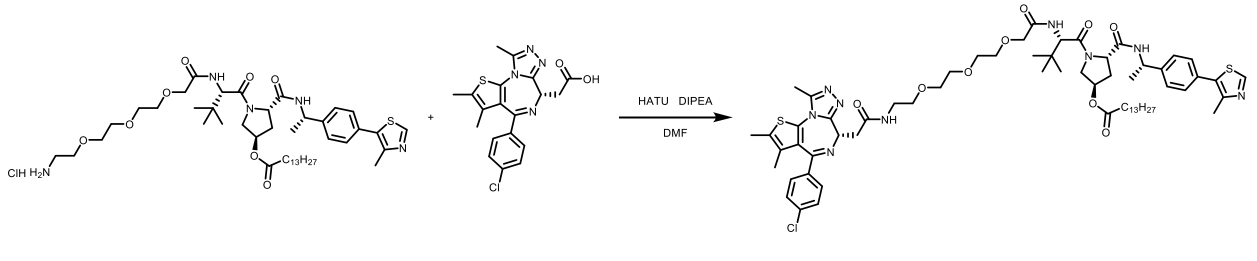


(3R,5S)-1-((S)-2-(tert-butyl)-17-((S)-4-(4-chlorophenyl)-2,3,9-trimethyl-6H-thieno[3,2-f][1,2,4]triazolo[4,3-a][1,4]diazepin-6-yl)-4,16-dioxo-6,9,12-trioxa-3,15-diazaheptadecanoyl)-5-(((S)-1-(4-(4-methylthiazol-5-yl)phenyl)ethyl)carbamoyl)pyrrolidin-3-yl tetradecanoate (26mg, 11% yield) was synthesized according to general procedure C. MS (ESI) m/z XYZ [M+H]$^1$H NMR (400 MHz, DMSO-D6) δ 8.98 (s, 1H), 8.47 (d, J = 7.7 Hz, 1H), 8.30 (t, J = 5.6 Hz, 1H), 7.47 (d, J

= 8.9 Hz, 2H), 7.42 (d, J = 3.1 Hz, 2H), 7.40 (q, J = 2.3 Hz, 2H), 7.37 – 7.34 (m, 2H), 5.17 (d, J = 17.7 Hz, 1H), 4.89 (p, J = 6.8 Hz, 1H), 4.51 – 4.33 (m, 3H), 3.93 (d, J = 4.7 Hz, 2H), 3.75 (dd, J = 11.7, 4.0 Hz, 1H), 3.62 – 3.55 (m, 8H), 3.30 – 3.18 (m, 4H), 2.58 (s, 3H), 2.44 (s, 3H), 2.39 (d, J = 0.9 Hz, 3H), 2.24 (td, J = 7.4, 2.5 Hz, 3H), 1.96 (s, 1H), 1.60 (d, J = 0.9 Hz, 3H), 1.46 (t, J = 8.4 Hz, 3H), 1.35 (d, J = 6.9 Hz, 3H), 1.21 (s, 22H), 0.93 (d, J = 6.9 Hz, 9H), 0.85 – 0.80 (m, 3H).

**(3R,5S)-1-((S)-2-(tert-butyl)-17-((S)-4-(4-chlorophenyl)-2,3,9-trimethyl-6H-thieno[3,2-f][1,2,4]triazolo[4,3-a][1,4]diazepin-6-yl)-4,16-dioxo-6,9,12-trioxa-3,15-diazaheptadecanoyl)-5-(((S)-1-(4-(4-methylthiazol-5-yl)phenyl)ethyl)carbamoyl)pyrrolidin-3-yl stearate (GAL-02-248)**

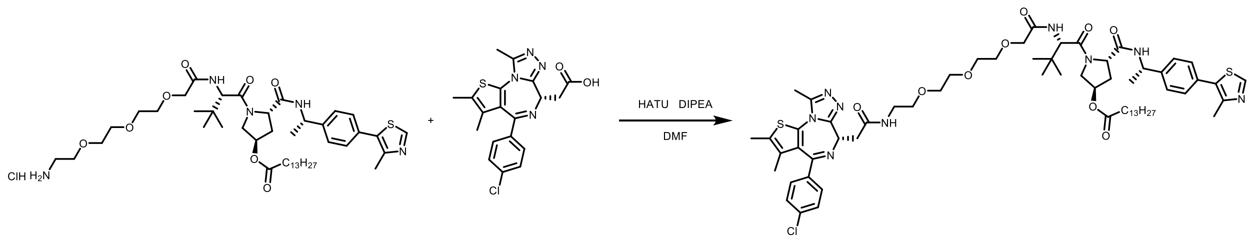


(3R,5S)-1-((S)-2-(tert-butyl)-17-((S)-4-(4-chlorophenyl)-2,3,9-trimethyl-6H-thieno[3,2-f][1,2,4]triazolo[4,3-a][1,4]diazepin-6-yl)-4,16-dioxo-6,9,12-trioxa-3,15-diazaheptadecanoyl)-5-(((S)-1-(4-(4-methylthiazol-5-yl)phenyl)ethyl)carbamoyl)pyrrolidin-3-yl stearate (18.7mg, 11% yield) was synthesized according to general procedure C. MS (ESI) 641 [M/2].

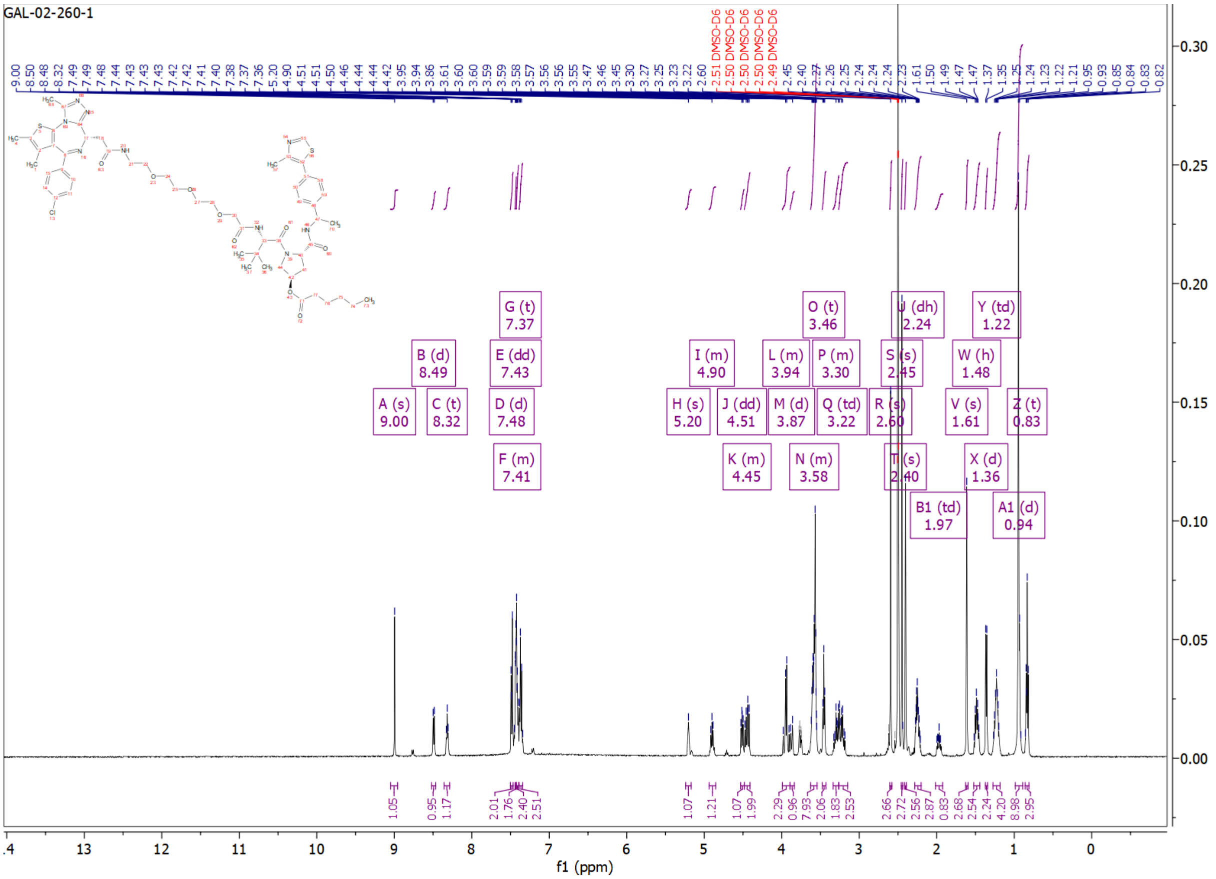
GAL-02-260-1
A (s) 9.00
B (d) 8.49
C (t) 8.32
D (d) 7.48
E (dd) 7.43
F (m) 7.41
G (t) 7.37
H (s) 5.20
I (m) 4.90
J (dd) 4.51
K (m) 4.45
L (m) 3.94
M (d) 3.87
N (m) 3.58
O (t) 3.46
P (m) 3.30
Q (td) 3.22
R (s) 2.60
S (s) 2.45
T (s) 2.40
U (dh) 2.24
V (s) 1.61
W (h) 1.48
X (d) 1.36
Y (td) 1.22
Z (t) 0.83
A1 (d) 0.94
B1 (td) 1.97
f1 (ppm)

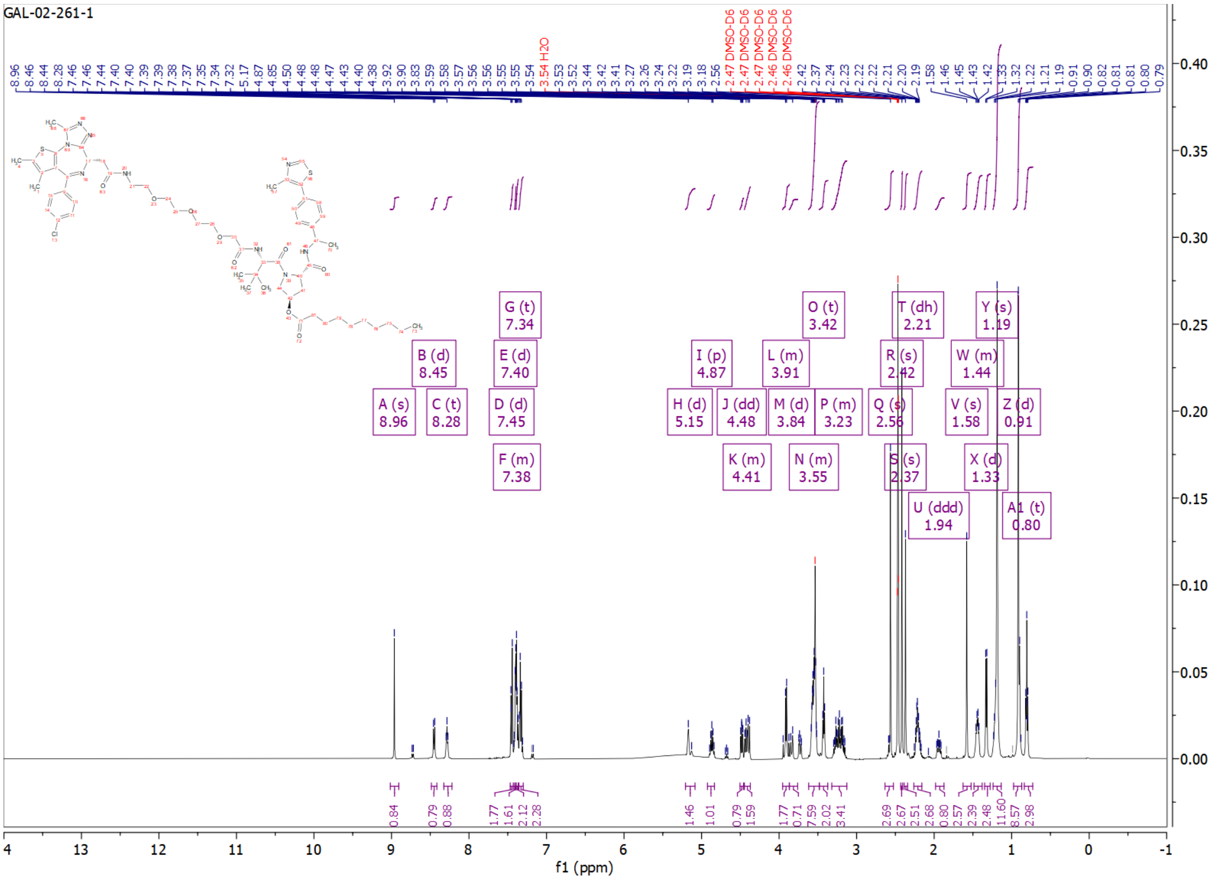
GAL-02-261-1
3.54 H2O
2.47 DMSO-D6
2.47 DMSO-D6
2.47 DMSO-D6
2.47 DMSO-D6
2.46 DMSO-D6
2.46 DMSO-D6
A (s) 8.96
B (d) 8.45
C (t) 8.28
D (d) 7.45
E (d) 7.40
F (m) 7.38
G (t) 7.34
H (d) 5.15
I (p) 4.87
J (dd) 4.48
K (m) 4.41
L (m) 3.91
M (d) 3.84
N (m) 3.55
O (t) 3.42
P (m) 3.23
Q (s) 2.56
R (s) 2.42
S (s) 2.37
T (dh) 2.21
U (ddd) 1.94
V (s) 1.58
W (m) 1.44
X (d) 1.33
Y (s) 1.19
Z (d) 0.91
A1 (t) 0.80
0.84
0.79
0.88
1.77
1.61
2.12
2.28
1.46
1.01
0.79
1.59
1.77
0.71
7.59
2.02
3.41
2.69
2.67
2.51
2.68
0.80
2.57
2.39
2.48
11.60
8.57
2.98
f1 (ppm)

GAL-02-253-1

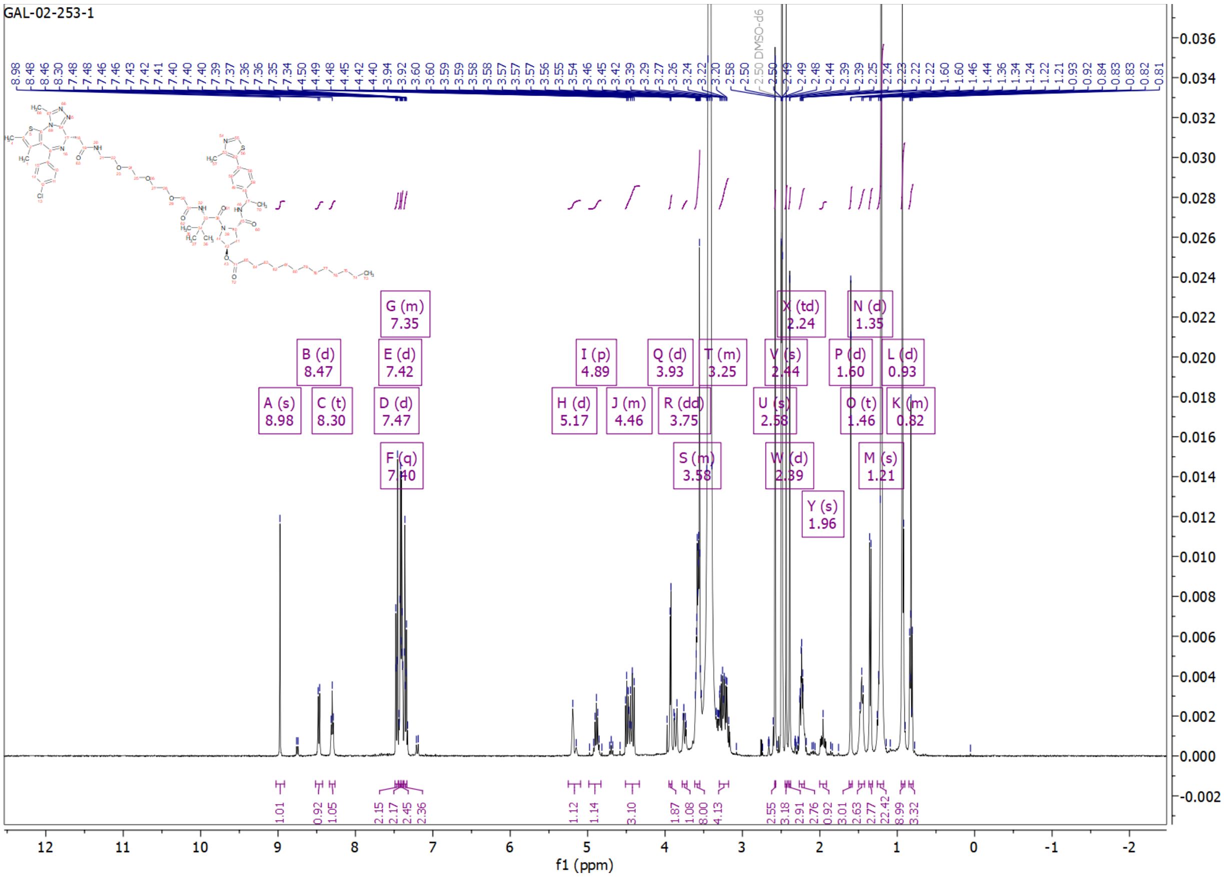

A (s) 8.98
B (d) 8.47
C (t) 8.30
D (d) 7.47
E (d) 7.42
F (q) 7.40
G (m) 7.35
H (d) 5.17
I (p) 4.89
J (m) 4.46
Q (d) 3.93
R (dd) 3.75
S (m) 3.58
T (m) 3.25
U (s) 2.58
V (s) 2.44
W (d) 2.39
X (td) 2.24
Y (s) 1.96
P (d) 1.60
O (t) 1.46
N (d) 1.35
M (s) 1.21
L (d) 0.93
K (m) 0.82
DMSO-d6
f1 (ppm)

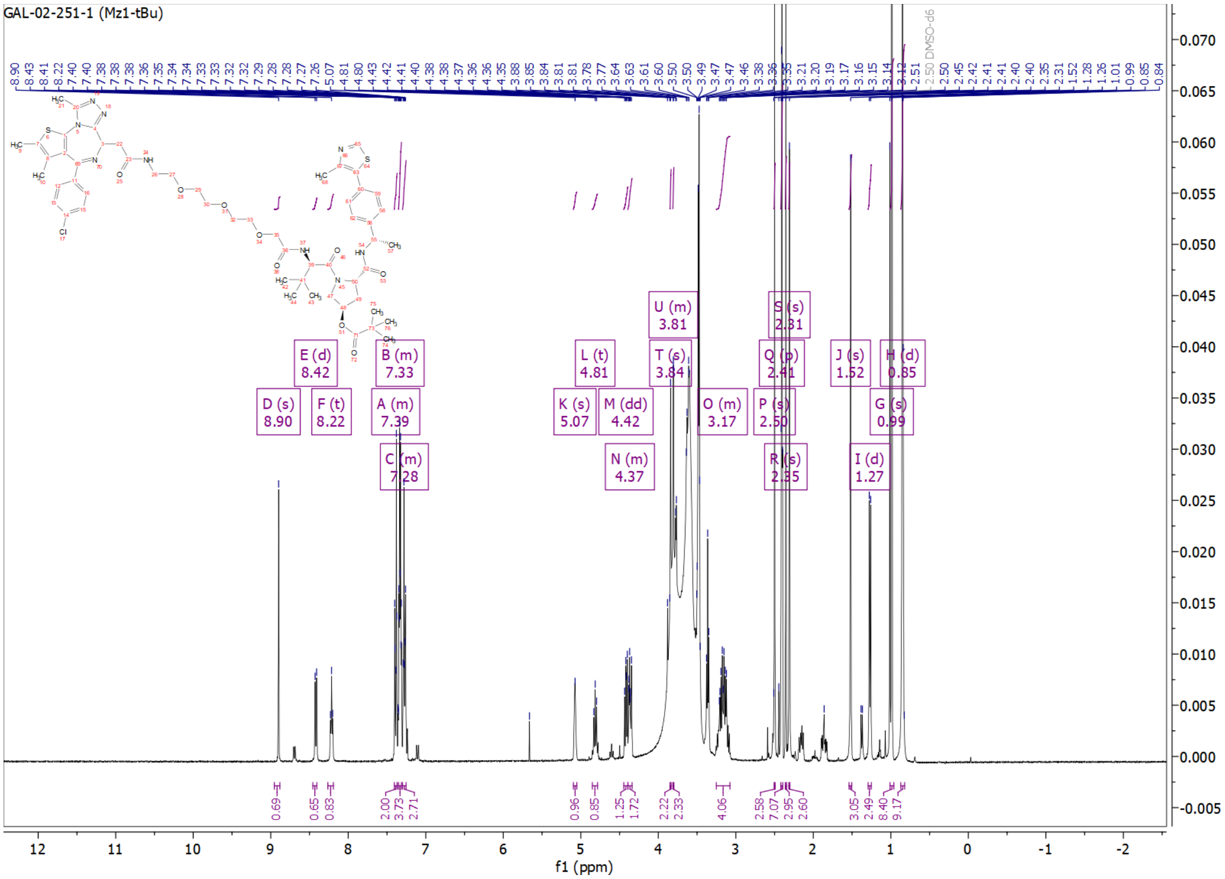
GAL-02-251-1 (Mz1-tBu)
D (s) 8.90
E (d) 8.42
F (t) 8.22
A (m) 7.39
B (m) 7.33
C (m) 7.28
K (s) 5.07
L (t) 4.81
M (dd) 4.42
N (m) 4.37
T (s) 3.84
U (m) 3.81
O (m) 3.17
P (s) 2.50
Q (p) 2.41
R (s) 2.35
S (s) 2.31
J (s) 1.52
I (d) 1.27
G (s) 0.99
H (d) 0.85
2.50 DMSO-d6
f1 (ppm)

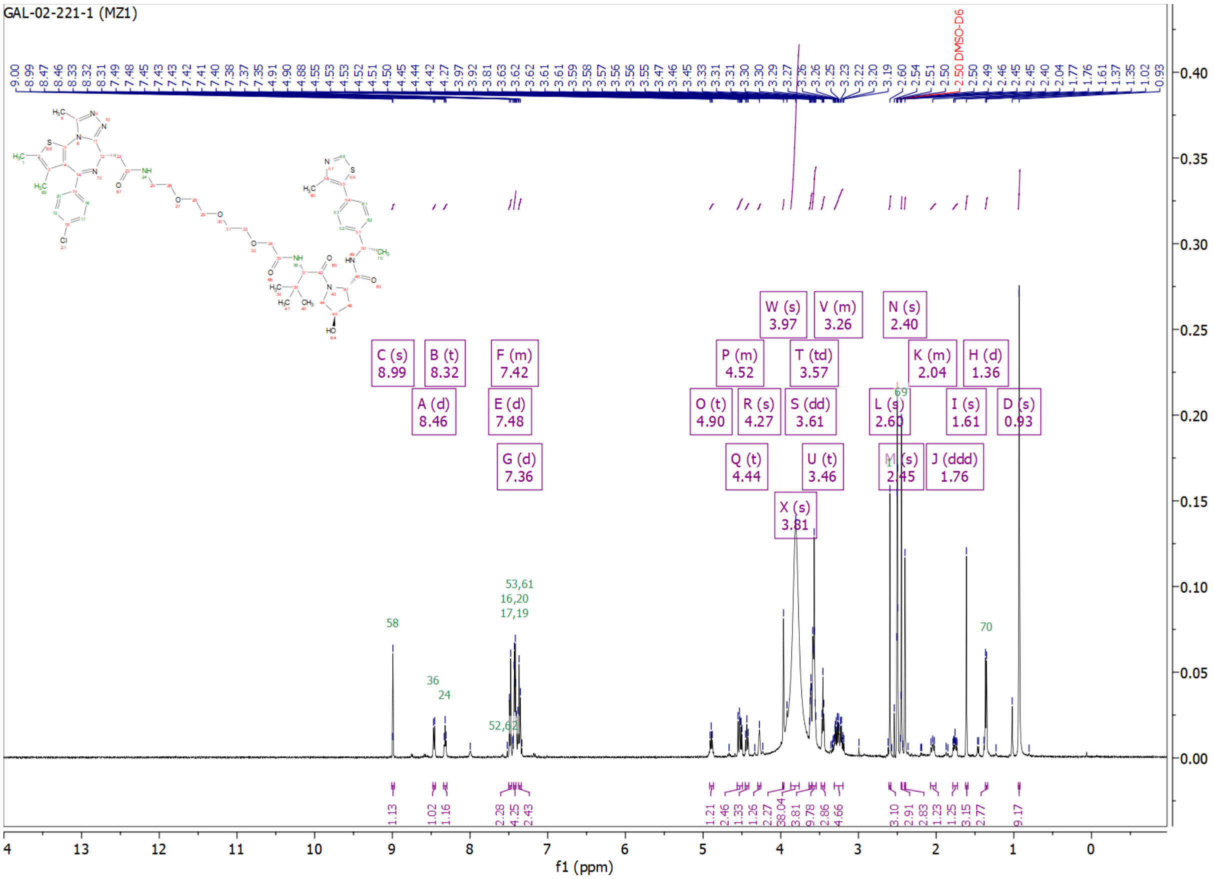
GAL-02-221-1 (MZ1)
2.50 DMSO-D6
C (s) 8.99
B (t) 8.32
A (d) 8.46
F (m) 7.42
E (d) 7.48
G (d) 7.36
W (s) 3.97
V (m) 3.26
P (m) 4.52
T (td) 3.57
O (t) 4.90
R (s) 4.27
S (dd) 3.61
Q (t) 4.44
U (t) 3.46
X (s) 3.81
N (s) 2.40
K (m) 2.04
H (d) 1.36
L (s) 2.60
I (s) 1.61
D (s) 0.93
M (s) 2.45
J (ddd) 1.76
f1 (ppm)

**Openlynx Report -** 

Sample: 55 Vial:2:13 ID:
File:GAL-02-221-1 Date:11-Jan-2024 Time:11:19:34
Description:MZ1

Printed: Thu Jan 11 11:25:38 2024

**Sample Report:**

**Sample 55 Vial 2:13 ID File GAL-02-221-1 Date 11-Jan-2024 Time 11:19:34 Description MZ1**

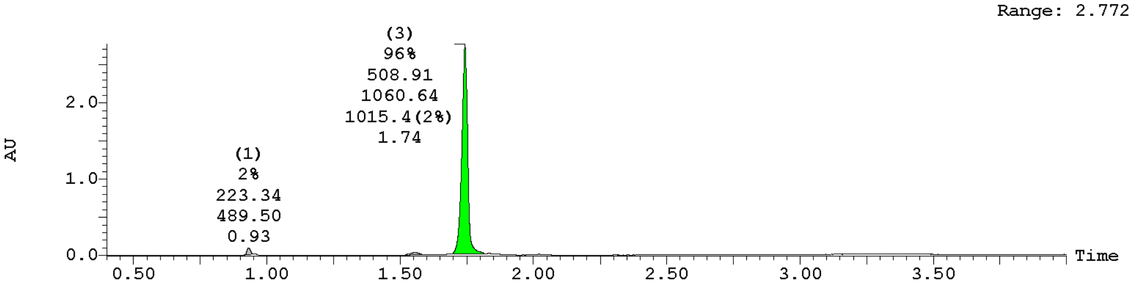


| Peak ID | Compound | Time | Mass Found |
|---|---|---|---|
| 1 | | 0.93 | Not Found |

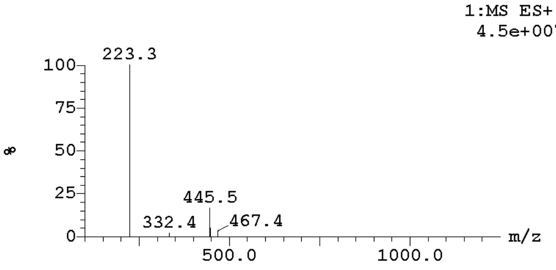


| Peak ID | Compound | Time | Mass Found |
|---|---|---|---|
| 2 | | 1.56 | Not Found |

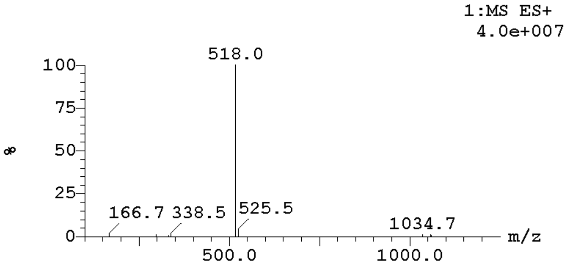


| Peak ID | Compound | Time | Mass Found |
|---|---|---|---|
| 3 | | 1.74 | Not Found |

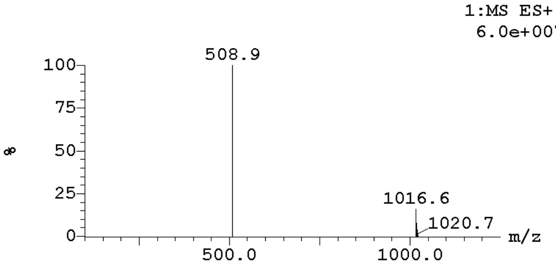


| Peak ID | Compound | Time | Mass Found |
|---|---|---|---|
| 1 | | 0.93 | Not Found |

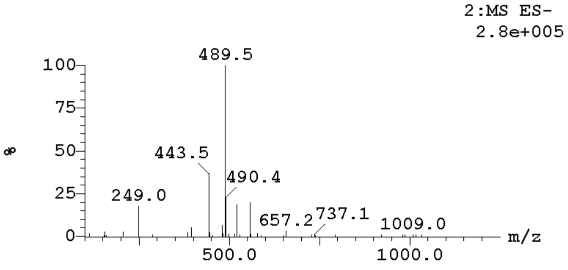

**Openlynx Report -**

Sample: 55 | Vial:2:13 | ID:
File:GAL-02-221-1 | Date:11-Jan-2024 | Time:11:19:34
Description:MZ1

Printed: Thu Jan 11 11:25:38 2024

**Sample Report (continued):**

| Peak ID | Compound | Time | Mass Found |
|---|---|---|---|
| 2 | | 1.56 | Not Found |

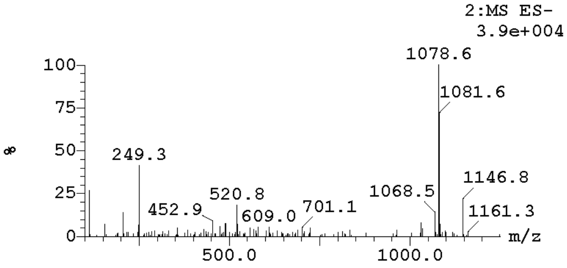


| Peak ID | Compound | Time | Mass Found |
|---|---|---|---|
| 3 | Found | 1.74 | 1015 |

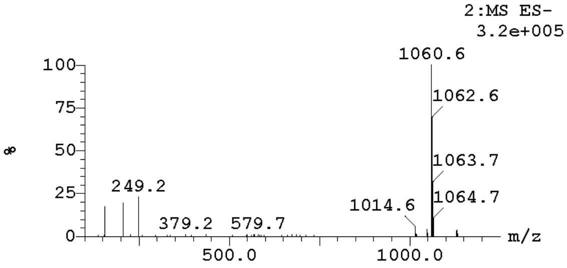

**Openlynx Report -** 

| | | |
|---|---|---|
| Sample: 47 | Vial:2:2 | ID: |
| File:GAL-02-251-1 | Date:08-Dec-2023 | Time:10:13:05 |
| Description:MZ1-tBu | | |

Printed: Fri Dec 08 10:31:00 2023

**Sample Report:**

**Sample 47 Vial 2:2 ID File GAL-02-251-1 Date 08-Dec-2023 Time 10:13:05 Description MZ1-tBu**

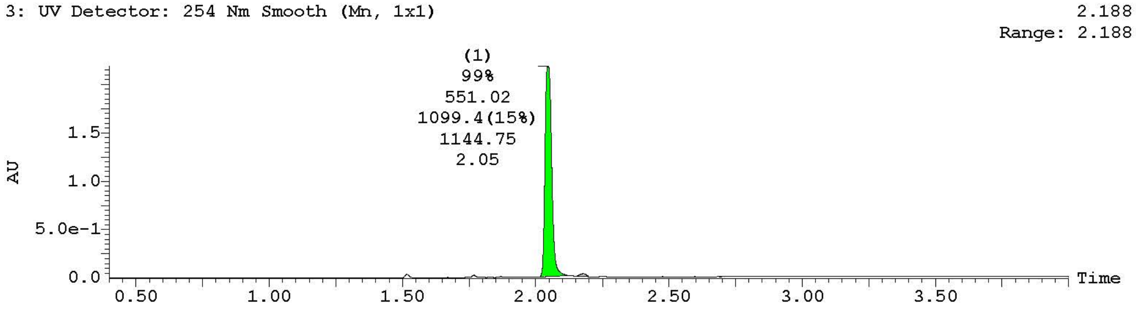


| Peak ID | Compound | Time | Mass Found |
|---|---|---|---|
| 1 | Tentative | 2.05 | 1101 |

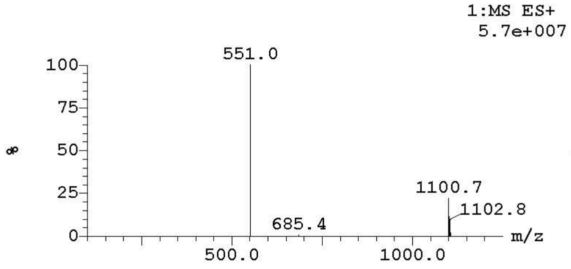


| Peak ID | Compound | Time | Mass Found |
|---|---|---|---|
| 2 | | 2.18 | Not Found |

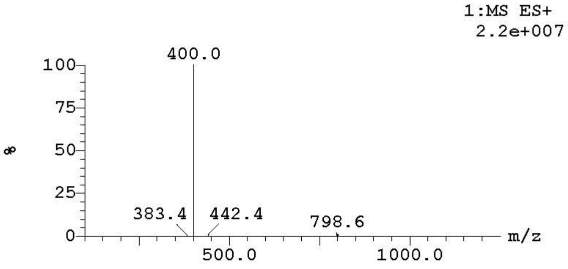


| Peak ID | Compound | Time | Mass Found |
|---|---|---|---|
| 1 | | 2.05 | Not Found |

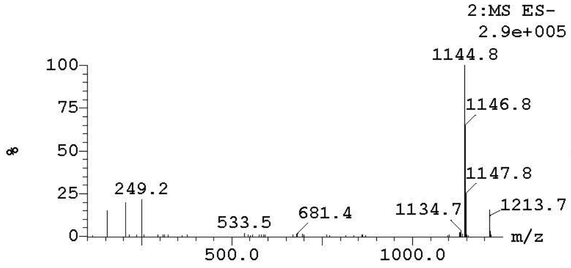


| Peak ID | Compound | Time | Mass Found |
|---|---|---|---|
| 2 | | 2.18 | Not Found |

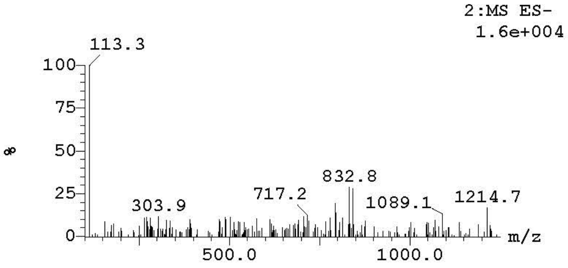

**Openlynx Report -** Page 1

| | | |
|---|---|---|
| Sample: 54 | Vial:2:8 | ID: |
| File:GAL-02-260-1-HPLC2 | Date:10-Jan-2024 | Time:10:06:14 |
| Description:MZ1-C6 (HPLC2/2) | | |

Printed: Thu Jan 11 11:27:19 2024

**Sample Report:**

**Sample 54 Vial 2:8 ID File GAL-02-260-1-HPLC2 Date 10-Jan-2024 Time 10:06:14 Description MZ1-C6 (HPLC2/2)**

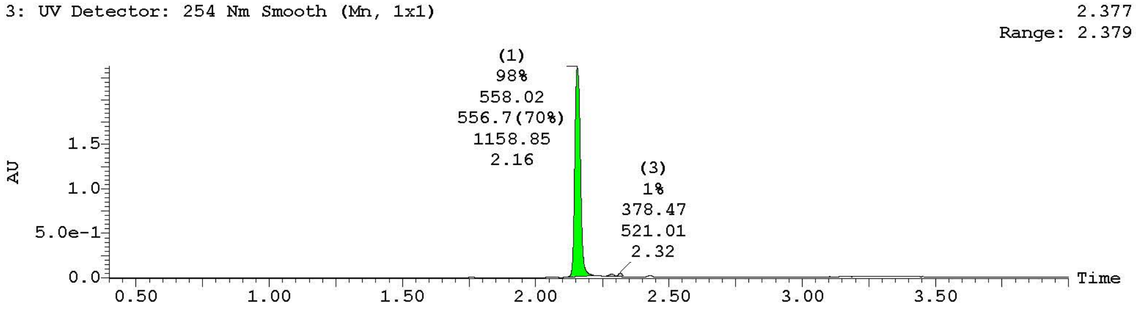


| Peak ID | Compound | Time | Mass Found |
|---|---|---|---|
| 1 | Found | 2.16 | 558 |

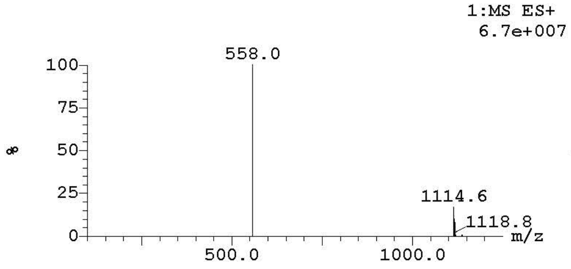


| Peak ID | Compound | Time | Mass Found |
|---|---|---|---|
| 2 | | 2.29 | Not Found |

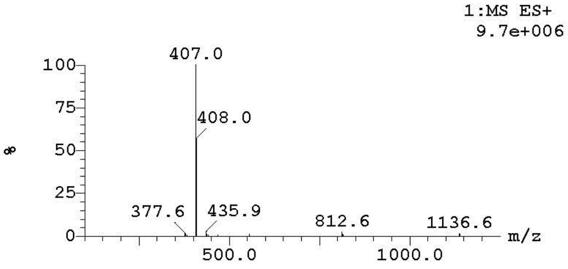


| Peak ID | Compound | Time | Mass Found |
|---|---|---|---|
| 3 | | 2.32 | Not Found |

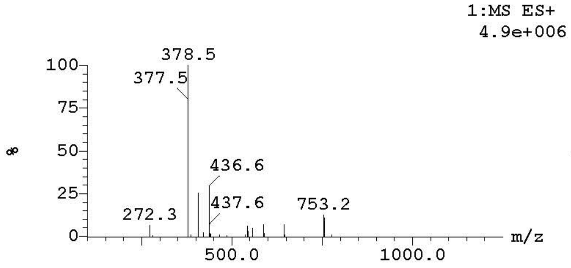


| Peak ID | Compound | Time | Mass Found |
|---|---|---|---|
| 1 | | 2.16 | Not Found |

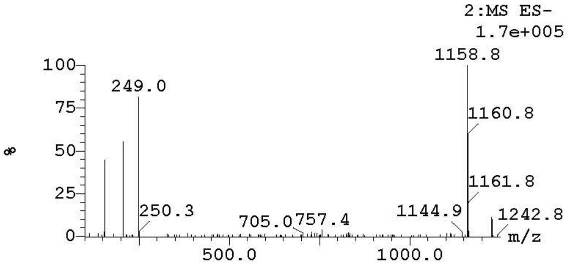

Sample: 54 Vial:2:8 ID:
File:GAL-02-260-1-HPLC2 Date:10-Jan-2024 Time:10:06:14
Description:MZ1-C6 (HPLC2/2)

Printed: Thu Jan 11 11:27:19 2024

**Sample Report (continued):**

| Peak ID | Compound | Time | Mass Found |
|---|---|---|---|
| 2 | | 2.29 | Not Found |

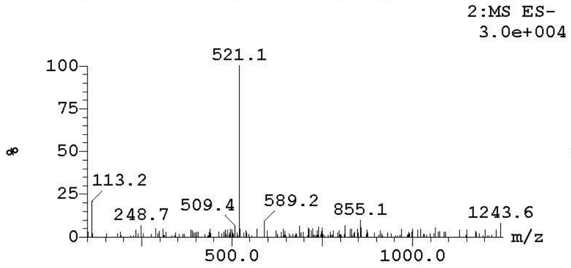


| Peak ID | Compound | Time | Mass Found |
|---|---|---|---|
| 3 | | 2.32 | Not Found |

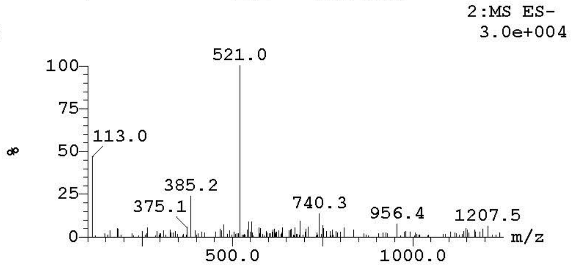

**Openlynx Report -** 

| | | |
|---|---|---|
| Sample: 53 | Vial:2:2 | ID: |
| File:GAL-02-261-1 | Date:08-Jan-2024 | Time:15:05:59 |
| Description:MZ1-C10 (HPLC1/3) | | |

Printed: Tue Jan 09 11:18:25 2024

**Sample Report:**

**Sample 53 Vial 2:2 ID File GAL-02-261-1 Date 08-Jan-2024 Time 15:05:59 Description MZ1-C10 (HPLC1/3)**

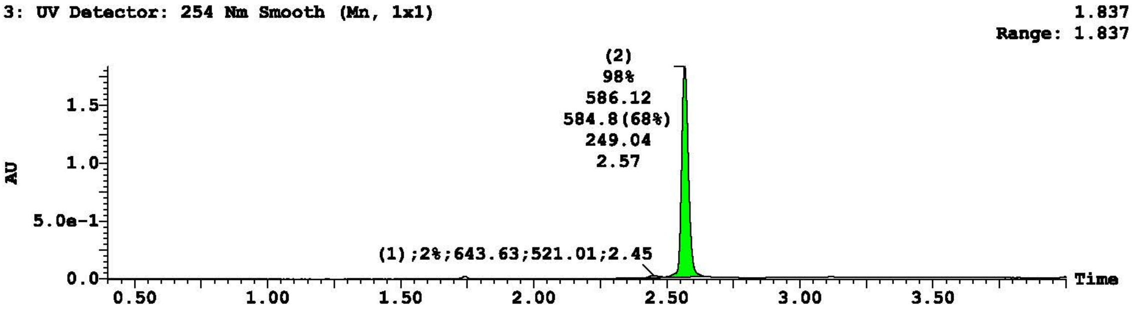


| Peak ID | Compound | Time | Mass Found |
|---|---|---|---|
| 1 | | 2.45 | Not Found |

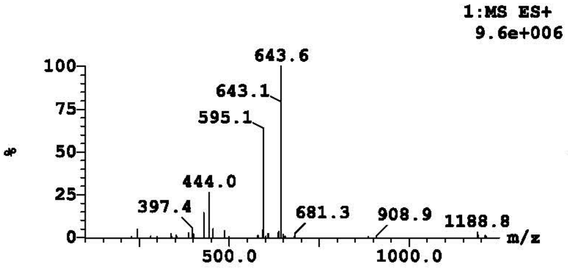


| Peak ID | Compound | Time | Mass Found |
|---|---|---|---|
| 2 | Found | 2.57 | 586 |

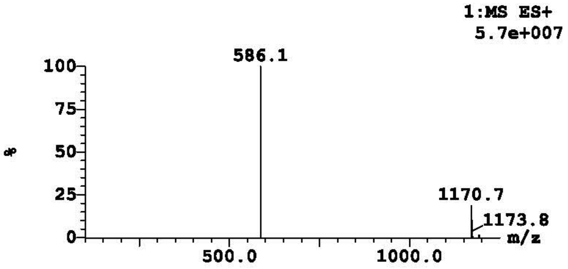


| Peak ID | Compound | Time | Mass Found |
|---|---|---|---|
| 1 | | 2.45 | Not Found |

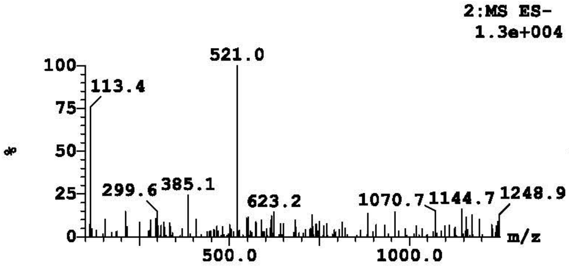


| Peak ID | Compound | Time | Mass Found |
|---|---|---|---|
| 2 | | 2.57 | Not Found |

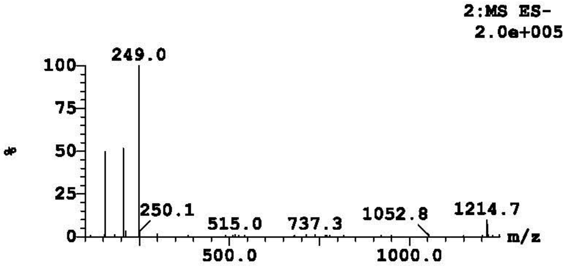

**Openlynx Report -**



Sample: 48 Vial:2:7 ID:
File:GAL-02-253-1 Date:13-Dec-2023 Time:09:36:33
Description:MZ1-C14

Printed: Wed Dec 13 11:13:15 2023

**Sample Report:**

**Sample 48 Vial 2:7 ID File GAL-02-253-1 Date 13-Dec-2023 Time 09:36:33 Description MZ1-C14**

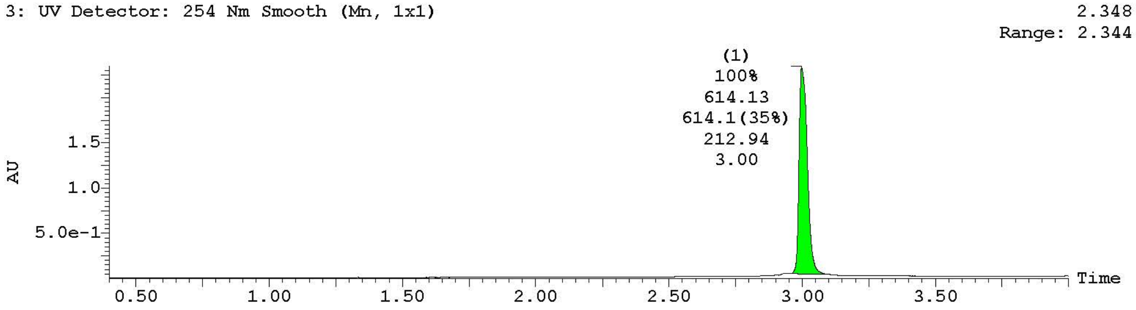


| Peak ID | Compound | Time | Mass Found |
|---|---|---|---|
| 1 | Found | 3.00 | 615 |

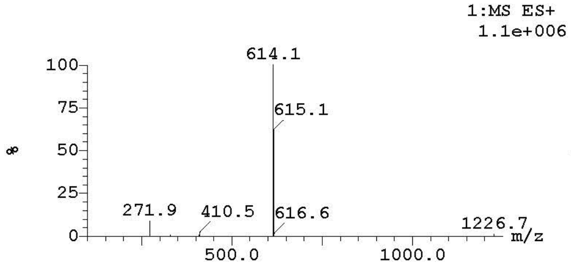


| Peak ID | Compound | Time | Mass Found |
|---|---|---|---|
| 1 | | 3.00 | Not Found |

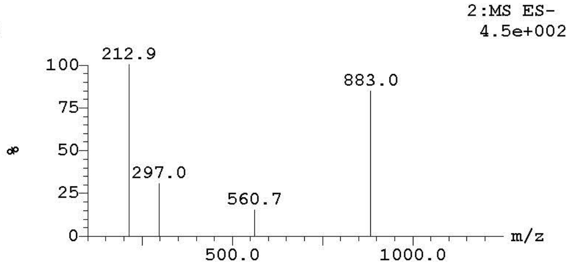

**Openlynx Report -**

| | | |
|---|---|---|
| Sample: 51 | Vial:2:11 | ID: |
| File:GAL-02-248-1 | Date:13-Dec-2023 | Time:14:43:28 |
| Description:MZ1-C18 | | |



Printed: Mon Dec 18 08:56:42 2023

**Sample Report:**

**Sample 51 Vial 2:11 ID File GAL-02-248-1 Date 13-Dec-2023 Time 14:43:28 Description MZ1-C18**

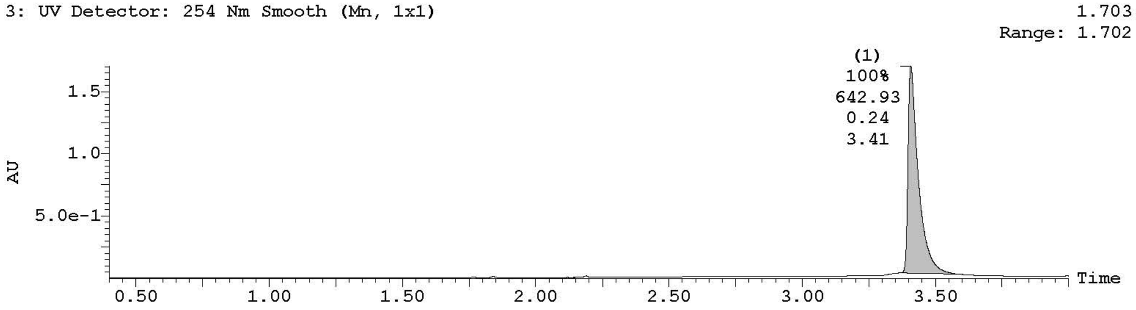


| Peak ID | Compound | Time | Mass Found |
|---|---|---|---|
| 1 | | 3.41 | Not Found |

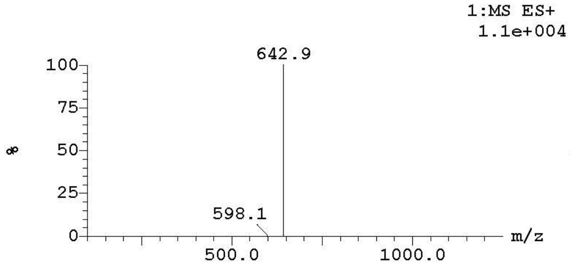


| Peak ID | Compound | Time | Mass Found |
|---|---|---|---|
| 1 | | 3.41 | Not Found |

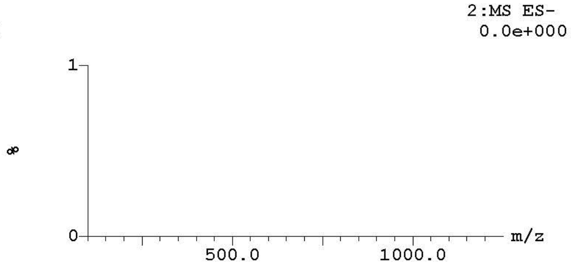
